# Insertions and deletions mediated functional divergence of Rossmann fold enzymes

**DOI:** 10.1101/2022.05.16.491946

**Authors:** Saacnicteh Toledo-Patiño, Stefano Pascarelli, Gen-ichiro Uechi, Paola Laurino

**Author notes:** This article is dedicated to the memory of Dan S. Tawfik.

## Abstract

Nucleobase-containing coenzymes are considered the relics of an early RNA-based world that preceded the emergence of protein domains. Despite the importance of coenzyme-protein synergisms, their emergence and evolution remain poorly understood. An excellent target to address this issue is the Rossman fold, the most catalytically diverse and abundant protein architecture in Nature. Here, we investigatedted the two largest Rossman lineages, namely the nicotinamide adenine dinucleotide phosphate (NAD(P))-binding and the S-adenosyl methionine (SAM)-dependent superfamilies. With the aim to identify the evolutionary changes that lead to a switch in coenzyme specificity on these superfamilies, we performed structural and sequence-based Hidden Markov Models to systematically search for key motifs in their coenzyme-binding pockets. Our analyses revealed how insertions and deletions (InDels) reshaped the ancient β1−loop−α1 coenzyme-binding structure of NAD(P) into the well-defined SAM-binding β1−loop−α1 structure. To prove this observation experimentally, we removed an InDel of three amino acids from the NAD(P) coenzyme pocket and solved the structure of the resulting mutant, revealing the characteristic features of the SAM-binding pocket. To confirm the binding to SAM, we performed isothermal titration calorimetry measurements, validating the successful coenzyme switch. Molecular dynamics simulations also corroborated the role of InDels in abolishing NAD-binding and acquiring SAM binding. Our results uncovered how Nature utilized insertions and deletions to switch coenzyme specificity, and in turn, functionalities between these superfamilies. This work also establishes how protein structures could have been recycled through the course of evolution to adopt different coenzymes and confer different chemistries.

**Significance Statement:** Cofactors are ubiquitous molecules necessary to drive about half of the enzymatic reactions in Nature. Among them, organic cofactors (coenzymes) that contain nucleotide moieties are believed to be relics of a hypothetical RNA world. Understanding coenzyme-binding transitions sheds light onto the emergence of the first enzymes and their chemical diversity. Rossmann enzymes bind to 7 out of 10 nucleotide coenzymes, representing an ideal target to study how different coenzyme specificities emerged and evolved. Here we demonstrated how insertions and deletions reshape coenzyme-specificity in Rossmann enzymes by retracing the emergence of the SAM-binding function from an NAD-binding ancestor. This work constitutes the first example of an evolutionary bridge between redox and methylation reactions, providing a new strategy to engineer coenzyme specificity.

## Introduction

Nearly 50% of enzymes require small, non-proteinogenic molecules, termed cofactors, to catalyze their otherwise unattainable reactions ^1^. Cofactors can be divided into inorganic ions and organic molecules, so-called coenzymes, which usually contain nucleotide moieties ^2^. Already in the 70s Harold White pointed out that nucleotide coenzymes play a major role in contemporary metabolism ^3,4^. Since then, it has been widely suggested that coenzymes may be the fossils of a nucleotide-based ancient metabolism prior to the emergence of the first proteins ^2,5^. In fact, in the beginning of the 80s, the first examples of ribozymes were discovered, strongly supporting White’s hypothesis ^6,7^. Further evidence was provided by the binding of several coenzymes to riboswitches ^8–10^ and the chemical resemblance of some coenzymes to RNA ^3^. In modern enzymes, coenzymes as well as inorganic cofactors, can be seen as the most catalytic units of all molecular architectures ^8^ Thus, given their importance, it is crucial to understand how their binding modes emerged within protein scaffolds. This understanding would ultimately allow us to switch cofactors and coenzymes specificities and therefore, to modify functions across protein families and superfamilies.

It is fundamental to understand how coenzyme-binding modes emerged within protein scaffolds and the Rossmann fold offers an excellent model system due to its unusual ability to bind multiple, chemically diverse coenzymes. Rossmann enzymes are the most catalytically diverse in Nature. They catalyze more than 300 different reactions ^11^, covering 38% of all known metabolic pathways. At a structural level, the Rossmann protein core is embedded in 20% of the total structures deposited in the PDB database ^12^. This core binds to seven out of ten nucleotide coenzymes ^13^, an ability that is extremely rare, since most of the protein architectures bind to a single coenzyme ^5^. Two major representatives among Rossmann proteins are the nicotinamide adenine dinucleotide (phosphate) (NAD(P))-, and the S-adenosyl methionine (SAM)-binding superfamilies. They are classified by the Structural Classification of Proteins (SCOP) database ^14^ in two distinct folds, c.2 and c.66, respectively. Similarly, the Evolutionary Classification of Protein Domain Structures (ECOD) database gathers them under different topology levels, but it groups them into the same homology category ^15^, claiming their common ancestry, this finding is also substantiated by recent studies ^16–19^. From a chemical perspective, the NAD(P)- and SAM-binding domains present different catalytic properties, while the first is predominantly involved in oxidoreductase reactions (EC 1), the second group acts as transferases (EC 2). Thus, how did two clearly homologous enzymes acquire their ability to bind different coenzymes, and as a result, distinct chemical activities?

Previous research has shown how semi-rational substitutions in Rossman enzymes resulted in a change of coenzyme specificity between NAD and NADP ^20,21^. However, NAD and NADP are closely related in structure, differing by only a phosphate moiety, and perform the same chemistry. Thus, this example does not provide information about how different chemistries evolved. In addition, the exploration of random mutations has demonstrated the important role of substitution in the emergence of new protein functions in general ^22^. In contrast, the role of insertions and deletions (InDels) in shaping enzymatic functions remains largely unexplored ^23^. Not only InDels are approximately 100 times more likely to have a deleterious effect ^24,25^ than substitutions, but also more difficult to explore in protein libraries since most approaches generate frame-shifting InDels at high frequency (> 66%) ^26^. Thus, there is a need to understand InDel-mediated changes in coenzyme specificity and protein evolution in general. This knowledge may have a great potential for enzyme engineering.

In this study, we systematically compared NAD(P)- and SAM-binding motifs of Rossmann enzymes, identifying well-defined and highly conserved β1−loop−α1 structural features for each coenzyme pocket (**Figure 1**). Despite the low sequence identity, assessment of sequence-based profile-profile comparisons revealed that InDels of up to 3 amino acids at the helix α1 link the structural features and coenzyme-binding motifs of these two superfamilies. Thus, we removed the three identified amino acids from two members of the NAD(P)-binding Rossmann superfamily and characterized the resulting mutants biophysically and biochemically. The X-ray structures of the mutants revealed the successful reshaping of the NAD(P)-binding β1−loop−α1 structure into the characteristic SAM-binding β1−loop−α1 structure. Similarly, binding measurements validated the coenzyme-binding switch from NAD to SAM. In addition, molecular dynamics simulations on the wild type and mutant scaffolds showed distinct hydrogen bonding networks, shaping the β1−loop−α1 motifs for each coenzyme pocket. The simulations showed how shortening the helix α1 disrupts a key interaction to the NAD-diphosphate group that ultimately abolishes NAD binding. Overall, these results show that a coenzyme-binding switch in Rossmann enzymes can be meditated by InDels, providing the first evidence of an InDel-based strategy employed by Nature to trade cofactors to evolve new functions.

**Figure 1.**
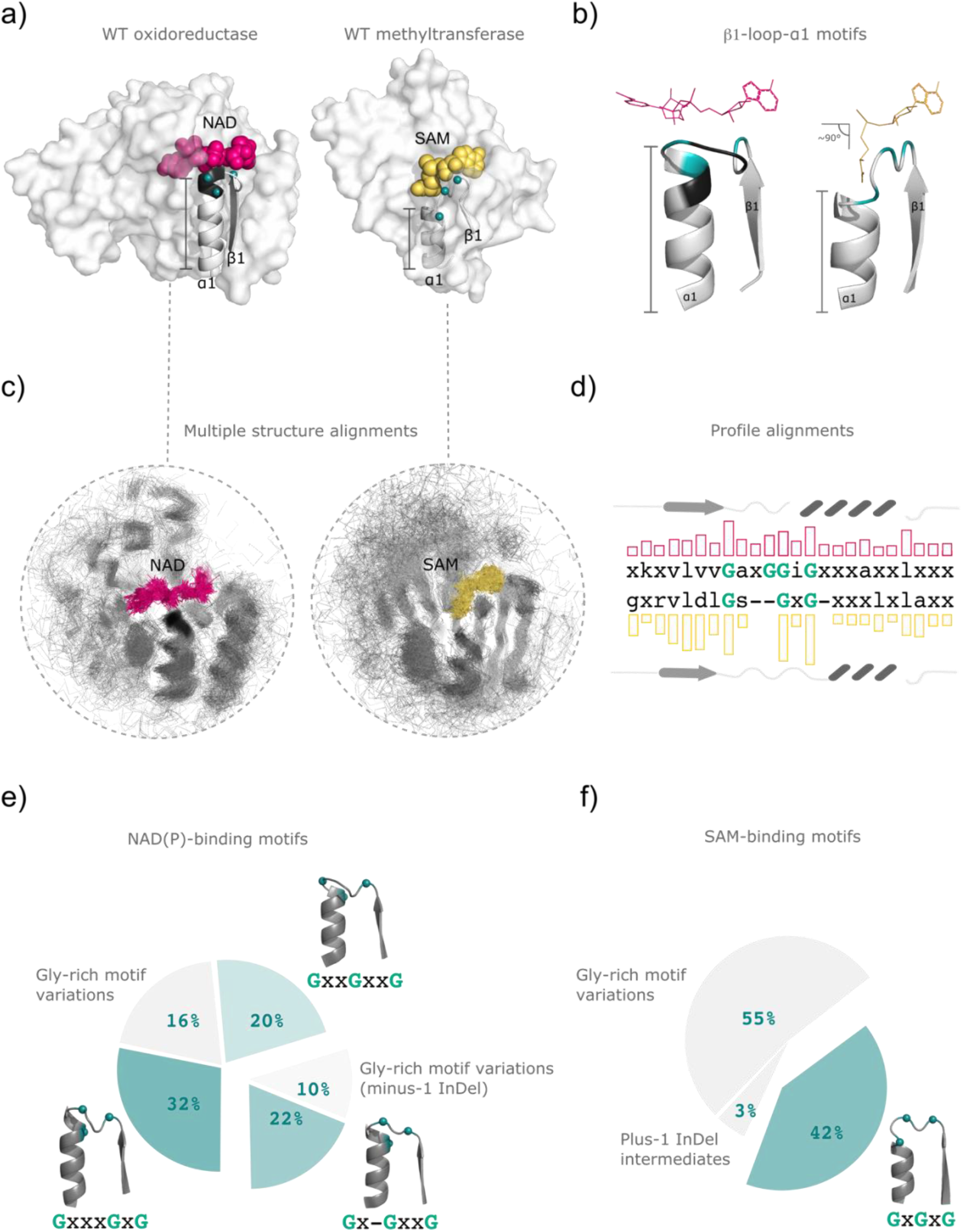
InDels within the coenzyme-binding pockets of NAD(P) and SAM-dependent Rossmann enzymes, mediate their specificity. (**a**) Protein structures of the target oxidoreductase (PDB 2D1Y) and methyltransferase (PDB ID 2AS0) are represented as surface and their β1−loop−α1 elements in cartoon. The coenzymes nicotinamide adenine dinucleotide (NAD) and S-adenosyl methionine (SAM) are highlighted as pink and yellow spheres, respectively. Glycine-rich motives are represented as turquoise spheres. Difference in length at the helix α1 is colored in black. (**b**) Zoom into the β1−loop−α1 structural elements of the NAD- and SAM-binding pockets shows the binding modes of their coenzymes and the position of the InDels (black) that change the localization of the Gly-rich motifs (turquoise). NAD (pink) and SAM (yellow) are shown as lines. The shorter helix α1 confers enough space to accommodate SAM in its catalytically productive conformation (**Figure S3**), namely bent in a ∼ 90° angle, in contrast to the extended conformation adopted by NAD(P). (**c**) Multiple structure superpositions of CATH homology groups of NAD(P)-binding 3.40.50.720 (left) (**Table S4**) and SAM-binding 4.40.50.150 (right) (**Table S5**) domains. Each domain is represented as thin ribbon line and the coenzymes as sticks. Both superfamilies show a high structural resemblance at their N-terminal regions (**Figure S2**) and a highly conserved conformation of the coenzymes. Superimpositions were downloaded from the CAHT ^1^ database (**d**) Sequence-base profile alignments of NAD-binding and SAM-binding superfamilies revealed the position of their Gly-rich motifs (turquoise) and the resulting InDels (gaps). (**e**) Three main Gly-rich motifs observed within the NAD(P)-binding superfamily are: GxGxxxG (32%), GxGxxG (22%) and GxxGxxG (20%). The second motif is structurally related to the β1−loop−α1 of flavin adenine dinucleotide (FAD)-binding Rossmann domains and presents a deletion (minus-1 InDel) of one amino acid with respect to the others. Gly-rich motif variations are observed for both: main-motif (16%) and minus-1 InDel intermediates (10%). (**f**) A single Gly-rich motif overrepresents the SAM-binding pocket, GxGxG (42%). The remaining pockets present diverse Gly-rich motif variations (55%) that lack one or more Gly-residues within the consensus. A few variants (3%) display plus-1 InDel intermediates (**Figure S7**).

## Results

### Drawing evolutionary bridges between the two largest Rossmann superfamilies

To identify the structural and sequence features that confer NAD(P) and SAM-binding specificity, we first performed a comprehensive analysis on sequences with known structures of Rossmann NAD(P)-dependent oxidoreductases and SAM-dependent methyltransferases. We found that despite the local homology at the N-terminus of these superfamilies (**Figure S1 and S2**), they display distinct and well-conserved structural features at their β1−loop−α1 coenzyme-binding regions. The length of the helix α1 is one turn shorter in the methyltransferases compared to the oxidoreductases (**Figure 1a**). This missing region provides the necessary space to embed the methionine moiety of SAM into the protein core and allows SAM to adopt a bent conformation that is necessary to catalyze the methyl transfer (**Figure 1b and S3**). In contrast, NAD(P) is bound in an extended conformation, where the longer helix α1 allows the interaction of the loop between β1 and α1 with the NAD(P)-diphosphate group. These distinct β1−loop−α1 features are highly conserved within each superfamily, as illustrated by their multiple structural alignments (**Figure 1c**). Similarly, at a sequence level, their coenzyme-binding motifs differ (**Figure 1d**). In fact, different Gly-rich motifs have been reported within different families ^27^, as well as consensus for each protein superfamily ^16,28–30^. Analysis based on hidden Markov models indicated three main Gly-rich motifs for the NAD(P)-binding superfamily (**Figure 1e**): V/IxGxxxGxG (32%), V/IxGxxGxxG (20%), and a shorter motif (minus-1 InDel), which presents an amino acid deletion V/IGxGxxG (22%). This shorter coenzyme-binding structure corresponds to the flavin adenine dinucleotide (FAD)-binding β1−loop−α1 motif. The remaining structures displayed different variations to the Gly-rich motifs with respect to both, the longer and the minus-1 InDel motifs (16% and 10%), respectively. Interestingly, ten outliers, with respect to the longer NAD(P)-binding motif, presented a three-residue insertion (plus-3 InDel). This insertion confers the β1−loop−α1 structure that is observed in coenzyme A-binding pockets. Here we focused on the V/IxGxxxGxG and V/IxGxxGxxG motifs, hereafter combined as V/IxGxxGGxG. The SAM-binding pocket showed the motif D/ExGxGxG (42%) and variations lacking one or more glycine residues (55%) of the consensus (**Figure 1f**). Among the outliers, intermediates presenting an amino acid insertion (plus-1 InDel) were found (3%) (**Figure 1f** and **S7**). NAD(P) and SAM-binding motifs are interconvertible through InDels, indicating that the removal of three residues from the NAD(P) β1−loop−α1 region could lead to the highly conserved structural features observed for the SAM β1−loop−α1 elements.

### Three deletions in the NAD-binding motif led to the structural features of the SAM pocket

To determine whether the β1−loop−α1 motif is sufficient to modulate coenzyme specificity in Rossman enzymes, we aimed to engineer a coenzyme switch from NAD to SAM by modifying only this region. To select the best protein targets for the coenzyme switch, we generated sequence-based profile alignments as reported in the methods. These profiles identified the first half of both superfamilies as the most similar (**Figure S5**). These results are in line with the observation that Rossmann fold enzymes bind the common adenosine moiety of the coenzymes in this region ^19^. The hits obtained from sequence-based profile alignments were ranked according to sequence identity (**Table S1**). We selected oxidoreductase sequences that showed relatively high sequence similarity to methyltransferases. I.e., those that displayed at least 30% sequence identity over ≥ 60 residues. This threshold was carefully selected base on the homology benchmarking described by Sander *et al*, who suggest sequence identity cutoffs for given alignment lengths ^31^. We selected seven potential design targets and attempted their protein expression in *E. coli*. Among them, we then selected the short chain dehydrogenase/reductase (scDH) from *Thermus thermophilus* (**Figure 2a**) for further characterization due to its high expression yields and thermal stability. This dehydrogenase has a sequence identity of 30% over 67 residues with a hypothetical methylase from *Thermoplasma acidophilum* (PDB 1NE2), and 32% over 66 amino acids with the PrmA-like methylase (PDB 2NXE) from the same organism.

**Figure 2.**
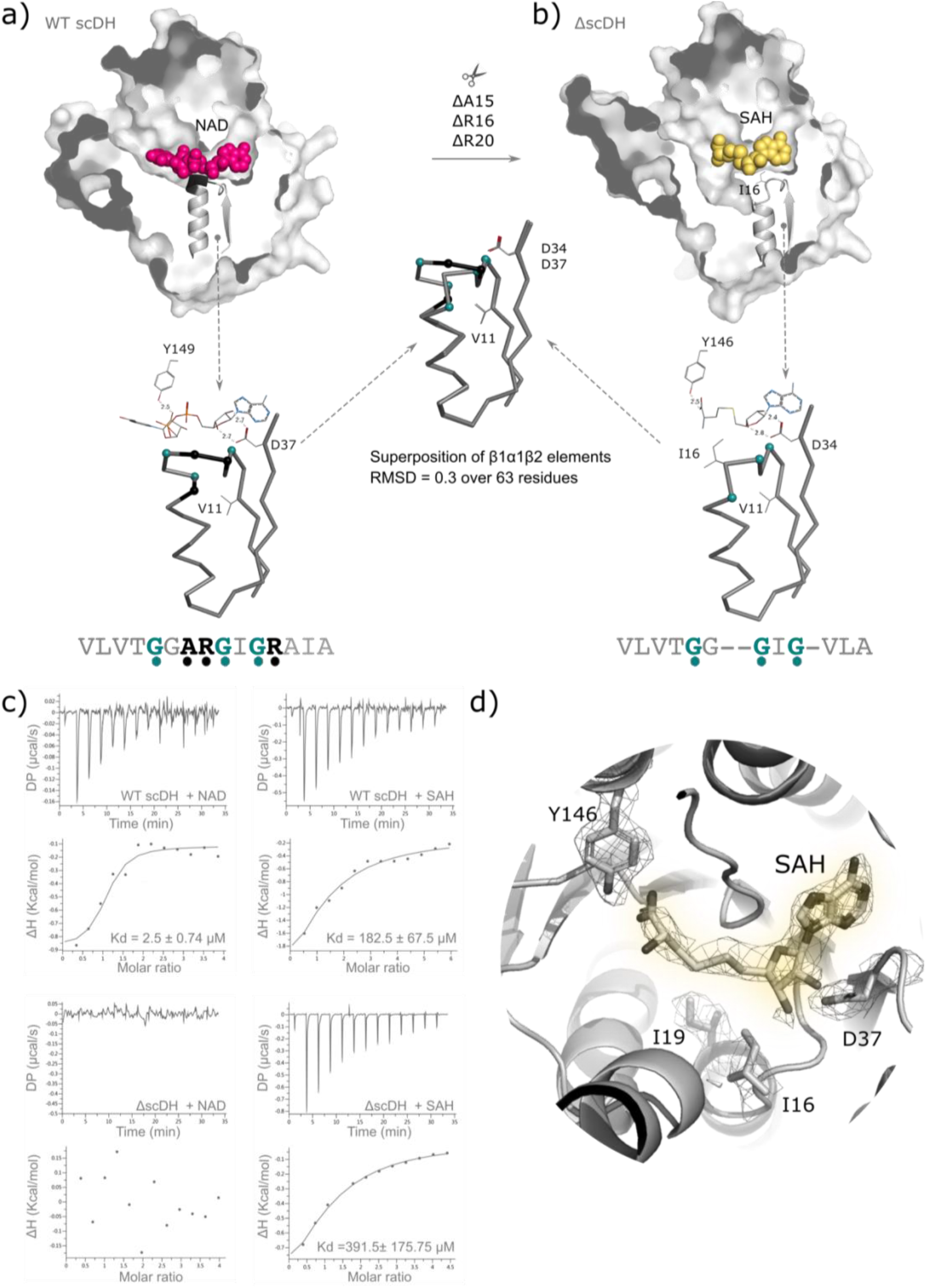
InDel extraction on the short chain dehydrogenase (scDH) from *Thermus thermophilus*. (**a**) Protein structures of the target protein scDH (PDB 2D1Y) and (**b**) the ΔscDH mutant (PDB 7XQM) are represented as surface and their β1−loop−α1 elements in cartoon. The coenzymes nicotinamide dinucleotide (NAD) and S-adenosyl methionine (SAM) are highlighted as pink and yellow spheres, respectively. The extraction of residues Ala15, Arg16 and Arg20 results in a shorter helix α1 in the ΔscDH mutant and opening of the NAD-binding pocket. Despite the resulting protuberance, SAH binds to the ΔscDH mutant in an extended conformation due to a blockage by Ile16 (gray lines). Zoom into the β1−loop−α1−β2 structural elements of the coenzyme pockets are displayed below their surface structures as ribbon and their structural alignment in the center. Removed residues are highlighted as black spheres and Gly-rich motifs as turquoise spheres. The respective coenzymes and their bidentate interaction of Asp37 (scDH) and Asp34 (ΔscDH) are shown as lines, highlighting a similar orientation of the adenosyl moiety. The interaction of Tyr149 (lines) with NAD in the WT scDH is also observed in the ΔscDH structure (Tyr 146) interacting with the amino acid moiety of SAH. Structural superposition of the β1−loop−α1−β2 reveals the successful shortening of the helix α1 and the typical structural motif of SAM-dependent methyltransferases (**Figure S4**). **(c)** Isothermal calorimetry titrations on the WT scDH revealed binding to NAD (Kd = 2.5 ± 0.74 μM) and an initial binding to SAH (Kd = 182.5 ± 67. μM). After the deletion, binding to NAD is abolished in the ΔscDH mutant, while the SAH binding remains (Kd = 391.5 ± 175.75 μM). (**d**) A zoom into the ΔscDH coenzyme pocket shows the electron density for: SAH, the obstructing residue Ile16, the Tyr146 interacting with SAH and the bidentate interaction of Asp37 to the ribosyl moiety. The ΔscDH structure has been solved at a 2.7 A resolution (R-free 0.18, R-work 0.18), the 2mFo-DFc map is contoured at 1.0 σ.

Three amino acids were removed from the coenzyme-binding pocket of NAD to generate the Gly-rich motif GxGxG of SAM. This was achieved by removing two amino acids (Ala15 and Arg16) between the first and third Gly, and one additional amino acid (Arg20) after the fourth Gly (VTGG<u>AR</u>GIG<u>R</u> ? VTGGGIG, **Figure 2a**). The deletions resulted in the mutant ΔscDH (**Figure 2b**), which expressed solubly and showed a melting point at 82 °C, similar to that at 85 °C observed for the wild type protein sdDH, reflecting the scaffold plasticity of Rossmann fold enzymes ^32^. To determine the coenzyme-binding affinities, isothermal titration calorimetry (ITC) experiments were conducted on the WT and ΔscDH mutant. S-adenosyl homocysteine (SAH) was chosen over SAM for the ITC measurements because of its chemical stability during the measurements. The WT protein scDH exhibited a K_d_ of 2.5 ± 0.7 μM for NAD. Surprisingly, scDH also showed binding to SAH with a K_d_ of 182.5 ± 67.5 μM. In contrast, after the deletions, the ΔscDH mutant lost the ability to bind NAD and exhibited a 2-fold lower affinity for SAH with a K_d_ of 392 ± 176 μM (**Figure 2c**).

To get insights into the binding mode of SAM, we solved the X-ray structure of the mutant ΔscDH in complex with SAH (**Table S2**). The refined holo structure revealed the desired shortened helix α1 and the β1−loop−α1 structural elements oriented similarly as in natural methyltransferases (**Figure 2b and S4**). However, SAH adopted an extended conformation instead of the catalytically competent bent conformation (**Figure 2d**), due to the interaction of its carboxy group with the hydroxy group of Tyr147, *via* hydrogen bonding. In natural methyltransferases, the SAM/SAH carboxy group mostly interacts with charged residues (mainly Asp/Glu) located within the β1 strand. However, in the ΔscDH mutant, this position is occupied by the hydrophobic residue, Val11 (**Figure 2b** and **S6**). Hydrophobic residues at this position are only observed for the spermidine synthase family (SCOP c.66.1.37), which catalyzes the transfer of a propylamine group to putrescine in the biosynthesis of spermidine ^33^. This family uses a decarboxylated form of SAM (dcSAM) as coenzyme, explaining why a charged residue at the beta strand β1 is not conserved. Looking at the β1−loop−α1 structural elements of spermidine synthases in more detail, revealed that this family displays a slight curvature at their coenzyme-binding loop, which closes the SAM binding pocket (**Figure S6**). A similar curvature was also observed for the ΔscDH mutant, whose residue Ile16 oriented towards the SAM-binding protuberance. To test the hypothesis that a charged residue is needed to induce SAM’s bent conformation, we introduced the substitutions Val11Glu, Val11Asp and Val11Asn in the wild type protein. Unfortunately, none of these mutations were tolerated, leading to insoluble protein expression of the resulting mutants.

### Coenzyme-switch on a malate dehydrogenase via InDels and one substitution

The above results on the ΔscDH mutant opened the following questions: are other oxidoreductases capable of binding SAM? While InDels are necessary to rebuild the SAM-binding β1−loop−α1 geometry and to abolish NAD binding, they are not sufficient for SAM to adopt the catalytically competent conformation. Thus, can this be achieved by introducing a charged residue at the beta strand β1? To address these questions, we selected the malate dehydrogenase (MDH) from *Escherichia coli* as the new scaffold (**Figure 3a**). This enzyme has been widely characterized and shows high expression yields in *E. coli*. Sequence-based profile alignments for MDH only showed that MDH shared relatively low probability of being related to SAM-binding methylases (HHsuite probabilities below 80%), being a top hit the O-methyltransferase (PDB 2QYO) from *Medicago truncatula* (**Figure S5**). This methylase aligns at 19% sequence identity over 93 residues. In analogy to the ΔscDH design, we removed three amino acids from MDH: two deletions (Ala9 and Gly10) between the first and third Gly; and a last one (Gln14) after the fourth Gly within the Gly-rich motif (GA<u>AG</u>GIG<u>Q</u> → GAGIG). In addition, we introduced three parallel substitutions: Val5Asp, Val5Glu and Val5Asn to induce the bent conformation of SAM. Since ΔMDH-V5N showed the highest expression, it was selected for further characterization. ITC measurements showed that the coenzyme switch was successful: The wildtype enzyme exhibited a K_d_ for NAD of 467 ± 148 μM and no initial binding for SAH, while the mutant ΔMDH-V5N completely abolished NAD binding and acquired 13.3 ± 7.2 μM affinity for SAH, providing evidence for the successful coenzyme-switch.

**Figure 3.**
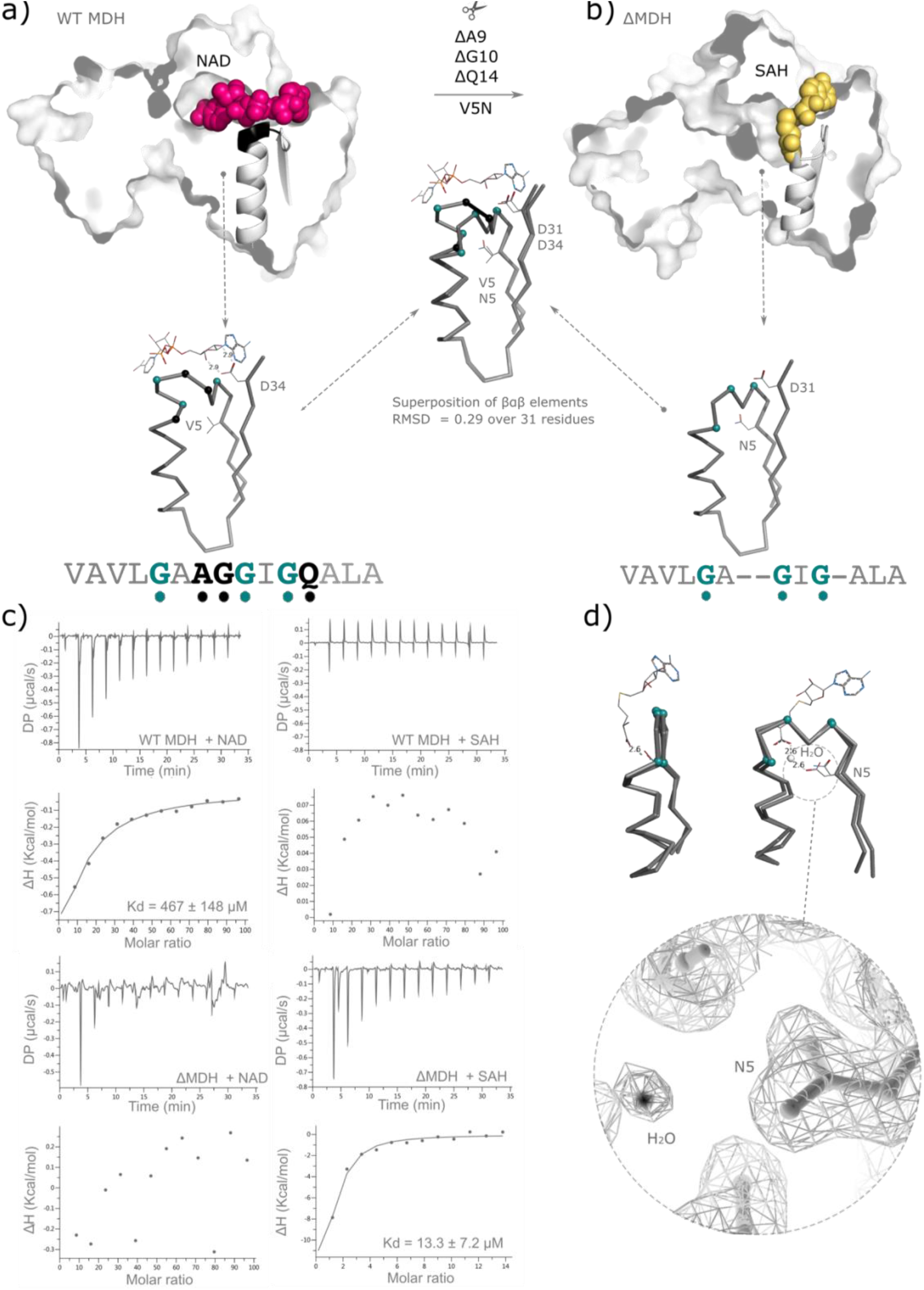
InDel engineering on malate dehydrogenase (MDH) reveals orthogonal coenzyme binding. (**a**) Protein structures of the target protein MDH (PDB 1EMD) and (**b**) the ΔMDH mutant (PDB 7XQN) are represented as surface and their β1−loop−α1 elements in cartoon. The coenzymes nicotinamide dinucleotide (NAD) and S-adenosyl methionine (SAM) are highlighted as pink and yellow spheres, respectively. The extraction of residues Ala9, Gly10 and Gln14 results in a shorter helix α1 in the ΔMDH mutant and an opening of the NAD-binding pocket towards the protein core. In contrast to the ΔscDH structure (**Figure 2b**), the coenzyme pocket is not obstructed, providing the necessary space to accommodate SAM in a bend conformation. SAM bent conformation was superimposed from the N5-glutamine methyltransferase from *Thermotoga maritima* (PDB 1NV8). Zoom into the β1−loop−α1−β2 structural elements of the coenzyme pockets are displayed below their surface structures as ribbon and their structural superposition in the center. Removed residues are highlighted as black spheres, Gly-rich motifs as turquoise spheres and coenzymes as lines. The bidentate interaction of Asp34 in MDH and the corresponding Asp31 residue in ΔMDH, are highlighted as lines. Structural superposition of the β1−loop−α1−β2 reveals the successful shortening of the helix α1 typical of SAM-dependent methyltransferase (**Figure S4**). The Val5Asn substitution (lines) shows the right orientation that allows the interaction to the amino acid moiety of SAM in SAM-dependent methyltransferases. **(c)** Binding studies via Isothermal titration calorimetry revealed that the wild type MDH binds to NAD (467 ± 148 μM) but not SAH, whereas the ΔMDH mutant loses the ability to bind NAD and acquires SAH binding (13.3 ± 7.2 μM), upon deletion. (**d**). Zoom into the substitution Val5Asn shows a water molecule (black sphere) interacting with Asn5 as in natural methyltransferases. The MDH structure has been solved at a 1.9 Å resolution (R-free 0.18, R-work 0.18) the 2mFo-DFc map in (d) is contoured at 1.0 σ.

To confirm the right loop region geometry and SAM conformation, we solved the structure of ΔMDH-V5N *via* X-ray crystallography. While the SAH cofactor could not be observed in the electron density, the apo structure revealed interesting features that suggested ΔMDH-V5N adopts the right loop conformation to accommodate SAM in its bent conformation. For instance, no curvature at the loop region was shown to close the SAM binding pocket as in the ΔscDH mutant and natural spermidine synthases (**Figure S6**). While the SAM binding pocket is partially blocked by Ile16 in ΔscDH, the pocket in ΔMDH is not obstructed (**Figure 2b** and **S6**). In fact, the obtained loop is flat and identical to those observed for natural methyltransferases (RMSD > 0.2). Finally, a water molecule interacts via H-bonding with the introduced Asn5. A similar interaction is observed to coordinate the SAM-carboxy group and the charged residues (Asp/Glu) in natural methyltransferases. Thus, ΔscDH-V5N recapitulates many of the structural features involved in coenzyme binding observed in native SAM-dependent methyltransferases.

### Molecular dynamics illustrate how InDels reshape H-bond networks across coenzymes pockets

We performed Molecular Dynamics (MD) simulations to further explain the molecular basis of the coenzyme-binding modes. We ran five repeats of 200 ns simulations of the tetrameric WT scDH (PDB ID: 2D1Y) without coenzyme (apo) or bound to four NAD molecules (holo), and the obtained crystal structure of the tetrameric ΔscDH mutant (PDB 7XQM) without coenzyme (apo) or bound to four SAH molecules (holo). The RMSD of all simulations is stable to a single peak (**Figure S9a-b**). In the ΔscDH, we observe an increased mobility of the β1-loop-α1 region nearby the deletion (positions 12-20), and a reduction of the mobility of the helix nearby the cofactor binding pocket (positions 180-210) (**Figure 4a**). Within the loop region, the increased mobility corresponded to a disruption of the hydrogen bonding network caused by the loss of two main interactions (Gly19-Ala23 and Ile18-Ile22) (**Figure 4b, c**). Interestingly, we also noticed that the WT scDH did not have a bond between the first and last glycine of the GxxxGxG motif (13Gly-19Gly). A bond in this position is considered important for loop formation in FAD/NAD(P) binding Rossman folds ^28^. In contrast, the helix in position 180-210 had smaller fluctuations in both apo and holo simulations of the ΔscDH mutant (**Figure 4a**), that might explain the increased sampling of conformations observed in the free energy surface (**Figure S9c**). Next, we looked at the hydrogen bonds between the protein and the coenzyme. We observed a higher number of bonds for NAD simulations, as expected by the difference in size with SAH (**Figure S9d**). For both proteins the most relevant residues that bind the coenzyme are those involved in binding the adenosine (**Figure 4d**). The main interactions are the bidentate contacts of Asp37 to the ribose, the backbone amino group of Leu58 to the adenine and the backbone carbonyl group of Ala86 to the ribose, and the respective positions in the deletion mutant (**Figure S10**). However, the top-ranking interaction involving Arg16 is missing in the mutant simulations. In the WT scDH, Arg16 interacts with the diphosphate group of NAD, which is absent in SAH. Further simulations of the WT scDH in complex with SAH confirmed that Arg16 does not take part in an important interaction with SAH (**Figure S9e**). In conclusion, the MD analysis shows that the delta mutant, although keeping most of the features of the WT, lacks an important residue for NAD binding that might be relatively superfluous in SAH binding.

**Figure 4.**
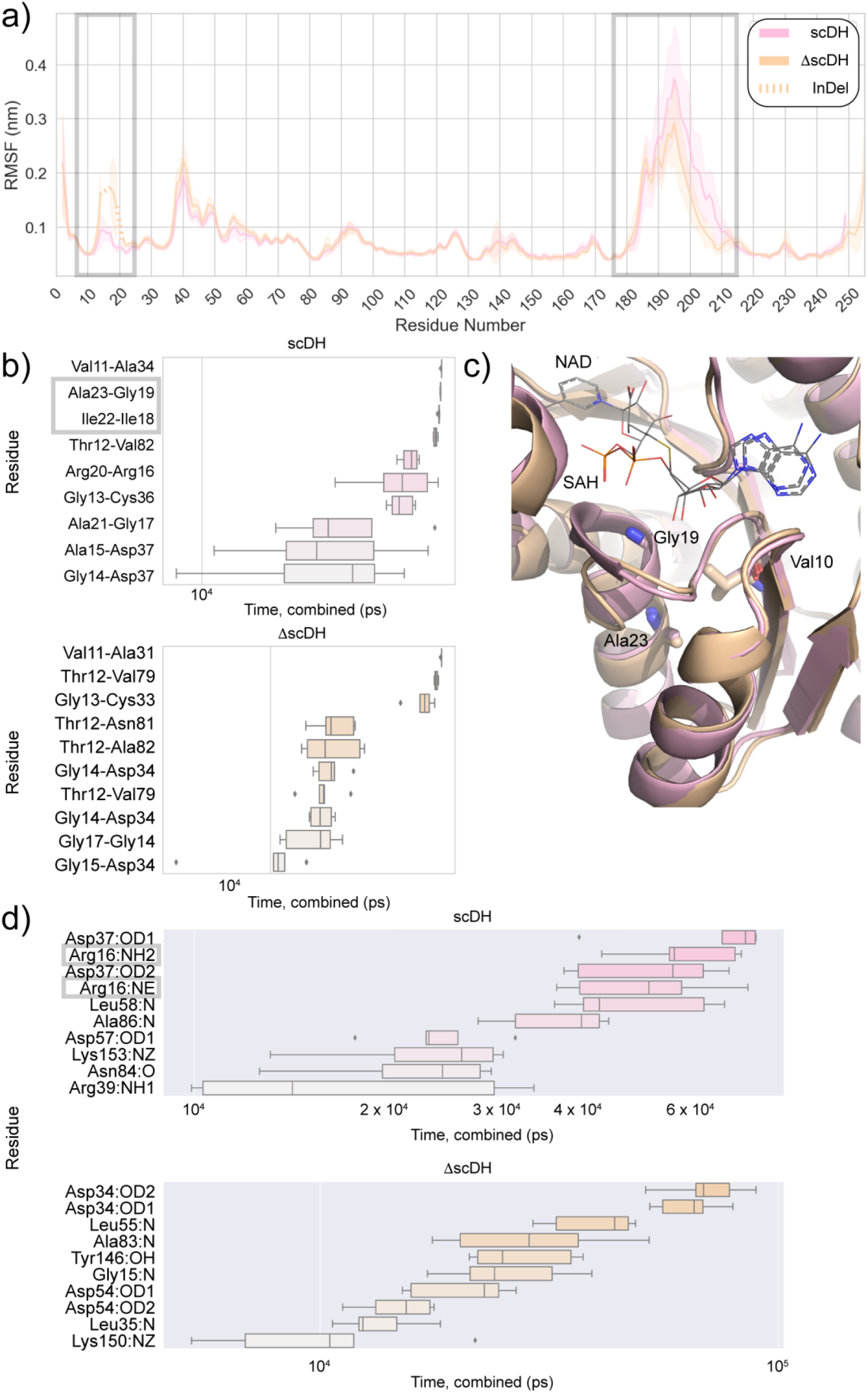
Molecular Dynamics analysis of coenzyme-binding on the short chain dehydrogenase and its delta mutant. **(a)** The root-mean-square-fluctuations (RMSFs) of the backbone atoms for the simulations of apo scDH and ΔscDH were compared. The X-axis shows the amino acid numbering of the wild type scDH as reference. The gray boxes highlight the differences discussed in the main text. The RMSFs of the wild type scDH are lower at the deleted region, but higher for residues Gly180 to Arg-200, from which Ile182 and Thr184 are in contact with the nicotinamide moiety, according to the holo crystal structure. **(b)** The box plots represent the prevalence time of a hydrogen bond during the simulations. Residues outside the β1−loop−α1 region (Leu10 to Ile22) have been omitted. Only the ten most prevalent hydrogen bonds are reported per simulation type. The gray boxes highlight two hydrogen bonds that are missing in the ΔscDH mutant. **(c)** A zoom into the coenzyme pockets of NAD and SAM. The structures of the wild type scDH (pink) and DscDH (yellow) are displayed in cartoon, NAD and SAM coenzymes as lines. The three stabilizing residues according to Kleiger and collogues ^2^ are highlighted as sticks. **(d)** The prevalence time of a protein atom hydrogen-bonding to the coenzyme. When multiple bonds are formed by the same atom in different moments, the time is accumulated. Gray boxes show the interactions of Arg16 to the NAD in the scDH simulations. This atom was deleted in the ΔscDH mutant. All simulations were performed in 5 repeats of 200 ns each. Knudsen, M. & Wiuf, C. The CATH database. *Hum. Genomics* **4**, 207 (2010). Kleiger, G. & Eisenberg, D. GXXXG and GXXXA motifs stabilize FAD and NAD(P)-binding rossmann folds through Cα-H…O hydrogen bonds and van der Waals interactions. *J. Mol. Biol*. **323**, 69–76 (2002).

## Discussion

This work addressed the unexplored role of InDels in the divergence of coenzyme-binding in Rossmann proteins. By introducing three InDels in the NAD binding pocket, we showed how this pocket can be remodeled to acquire the structural features of methyltransferases that allow SAM-binding in its catalytically productive bent-conformation.

### InDels drive coenzyme-binding and evolutionary trajectories in Rossmann enzymes

This work focused on switching coenzyme specificity from NAD to SAM. Although coenzyme engineering has been widely explored to invert the binding specificity of NAD and NADP ^20,21^ this approach differs to previous semi-rational mutagenesis approaches in the following aspects. It constitutes a unique example, where the switch involves coenzymes that catalyze fundamentally different functions, namely methylation and oxidoreductases reactions. For the first time, InDels are employed to reshape coenzyme pockets. A shorter NAD-binding helix α1 is essential to provide the necessary space for SAM to adopt the bent conformation (**Figure S3**). Finally, while previous switches between NAD and NADP mainly focus on the phosphate interaction, which is missing in NAD compared to NADP. Our InDel strategy remodeled hydrogen-bonding networks required for the well-defined β1−loop−α1 structures of the coenzyme pockets. Given the deleterious nature of InDels ^24^, it is remarkable that not only the binding loop, but also structural elements such as the helix α1 can be reconfigured, without disrupting the overall protein architecture and folding. This corroborates the scaffold plasticity observed for the Rossmann fold ^32^, which is believed to be one reason for its abundance, antiquity and its extraordinary multi-coenzyme-binding capabilities.

### An ancient phosphate binding loop as template for coenzyme-binding divergence

Common ancestry between two ancient Gly-rich motifs: the P-loop of ATPases and the phosphate-binding motif of Rossmann proteins has been recently suggested ^34^. Here, in addition, the fact that the NAD(P)-binding pocket was remodeled into the SAM-binding motif through an InDel extraction, suggests that other Gly-rich motifs may be interconvertible via InDels. A hint for this hypothesis is the existence of natural intermediates presenting InDels of different lengths (**Figure S7**). For instance, intermediates presenting minus-1 InDels within the β1−loop−α1 motif can be found in members of the NAD(P)-binding superfamily, this structural elements correspond to the β1−loop−α1 of flavin adenine dinucleotide (FAD)-binding domains, whose sequence similarity to NAD(P)-binders can be detected with HHsuite ^35^. Similarly, SAM-binding outliers display plus-1 InDels with respect to the most abundant motif that is discussed in this work (**Figure S4 and S7**). In addition, plus-3 InDels are observed for NAD(P)-binding motifs, which correspond in structure to the coenzyme A-binding pocket and show also sequence similarity, according to HHsuite ^35^ profiles. Although common ancestry may remain an unsolved problem, this InDel ladder strongly speaks against convergent evolution, showing how InDels may have finetuned different coenzyme pockets. This observation uncovers a potential strategy used by Nature to evolve new functions on similar protein structures, which can be reproduced experimentally *via* InDel engineering.

### Molecular Dynamics simulations revealed distinct H-bonding networks for the NAD and SAM coenzyme pockets

In line with the experimental data, the molecular dynamic (MD) simulations show that coenzyme specificity is determined by the InDels, given that the simulation is ergodic and sufficiently long. The shortening of the helix α1 increases the mobility of the resulting β1-loop-α1 structure, disrupting a key interaction with NAD(P) and opening the coenzyme pocket. This interaction loss explains why NAD(P)-binding motifs do not present deletions larger that 2 residues at the helix α1, where NAD stretches and interacts. Larger InDels may be overrepresented in methyltransferases since the protuberance created towards the protein core is needed to accommodate SAM in its catalytically productive conformation. The NAD(P)-binding Rossmann superfamily displays a Gly-rich motif that is similar to the FAD/NAD(P)-binding superfamily ^36^. However, a plus-1 InDel in the coenzyme-binding motif causes a rearrangement of the Gly-residues and a distinct structure. Additionally, the H-bond characteristic of the GxGxxG motif between the first and last Gly was not observed in the scDH MD simulations. The variation of the Gly-rich motifs modulated by InDels hints to the plasticity and adaptability of Rossmann folds ^32^.

In agreement to a previous report on NAD(P)/FAD-binding Rossmann folds ^28^, we found that the WT scDH presents an additional Gly-rich motif (GxxxG/A) adjacent to the main. This motif also contributes to stabilize the β1-loop-α1 structural elements. The Gly/Ala residue of the motif formed one of the most persistent H-bond interactions (Gly19-Ala23), helping the α1 helix formation. This interaction acts in unison with the beta-sheet-forming H-bonds between Val11-Ala34 and Gly13-Cys36. Together with Ile18-Ile22, the Gly-Ala H-bond maintain the structural features required to bind NAD. Consistently, when these hydrogen bonds are missing in the ΔscDH mutant, the ability to bind NAD is lost. However, the SAM binding is not affected by this change. The MD data and structural analysis demonstrated how InDels in the β1-loop-α1 region of Rossmann enzymes had a minimum impact on the overall structure, while having maximum local effect on the coenzyme pockets, conferring distinct specificities.

### Ligand conformations provide hints on the evolutionary trajectory of Rossmann proteins

The fact that oxidoreductases can bind to SAM/SAH in an extended conformation further supports the existence of a promiscuous Rossmann ancestor. In fact, the case of the scDH presented here is not an isolated case. The structure of a previously reported putative Zn-dependent alcohol dehydrogenase (PDB 3IV6), displays intermediate structural features between NAD(P) and SAM-binding pockets (**Figure S8**). This oxidoreductase has been crystallized in complex with SAM in an extended conformation similar to that of dcSAM in aminopropyl transferases, and the SAH in the ΔscDH structure. Following the hypothesis of coenzymes being catalytic relics that existed prior to the emergence of proteins and considering the strict coenzyme geometries needed for catalysis, a malleable coenzyme pocket presents a plausible strategy to incorporate proteins envelopes around existing coenzyme conformations instead of creating protein scaffolds from scratch. In fact, it has been suggested that although coenzyme redundancy may not be essential for metabolism, it represents a potential strategy that enabled the efficient usage of early enzymes ^37^. Thus, a moldable Rossmann fold motif may have served as a template, which evolved with the help of InDels and mutations. For example, the NAD(P)-binding motif could have been reshaped into the dcSAM pocket of aminopropyl transferases via InDels (**Figure S6**). Similarly, the dcSAM pocket could have evolved into a SAM-binding pocket with the help of a charged residue, to stabilize the bent conformation of SAM that allows methylation instead of aminopropyl transfer. However, although aminopropyl transferases appear to be an intermediate between NAD(P)-binding oxidoreductases and SAM-dependent methylases (**Figure S6**), an evolutionary directionality cannot be concluded from our findings.

## Materials and Methods

### Sequence-based profile-profile comparisons with HHsuite

To generate sequence-based Hidden Markov Models, multiple sequence alignments were built for SCOP ^14^ folds c.2 and c.66 contained in the astral database release 2.07 ^38^. In addition, multiple sequence alignments were generated for ECOD ^15^ homology groups 2003.1.1 and 2003.1.5. The alignments were generated employing PSI-BLAST ^39^, which is a program included in the build.pl protocol described by Söding and colleges ^35^. We employed this protocol to generate profile-profile comparisons with default parameters. The secondary structure prediction was turned off (ssm=0) in order to perform a strictly sequence-based search. The gathered homologous regions were aligned with TM-align ^40^ and manual selections with PDBeFold ^41^. The obtained alignments were filtered according to sequence identity and well-superimposed structures.

### Profile-profile structural comparisons with PROMALS

Gly-rich motifs for Rossmann oxidoreductase and methylase superfamilies were obtained through profile alignments with PROMALS ^42^. In total, 332 sequences were gathered for the NAD(P)-binding domain (c.2) and 144 sequences for SAM-binding domain (c.66) of SCOP ^14^ released 2.07 ^38^ at a cutoff of 40%. Similarly, we created a sequence subsets for the NAD-binding domains (2003.1.1, 477 sequences) and SAM-binding domains (2003.1.5, 291 sequences) according to ECOD ^15^ database version 20220113 at a cutoff of 40%. SCOP was selected for its well-known manual curation, whereas ECOD was selected for containing more sequences. Similar consensus sequences were obtained for each database.

### Structural superpositions

Structural alignments for CATH superfamilies 3.40.50.720 and 3.40.50.150 (**Table S4 and S5**) were downloaded from CATH database and edited with PyMoL to generate the images.

### Gly-rich motif sorting

To obtain the different Gly-rich motifs, domain coordinates corresponding to the SCOP database release 2.08 at 40% sequence identity were compared. To ensure the proper localization of the Gly-rich motifs, a manual inspection and superposition of the β1−loop−α1 structural motifs was performed for each fold C.2 and C.66, respectively.

### Cloning expression and purification

The wildtype scDH gene was synthesized by TWIST Bioscience (codon-optimized for expression in *E. coli*). PCR amplification was performed with primers 5’-CGGCCTGGTGCCGCGC and 3’-CGCAAGCTTGTCGACGGAGCTCGAATTC, both containing appropriated overlapping regions for INFusion® cloning (TaKaRa Bio Inc) into the target vector pET28a(+). This vector encodes for an N-terminal hexa-histidine affinity tag and confers Kanamycin resistance. The MDH gene was contained in the pASK-IBA3(+)-MDH vector, which has been kindly provided by Dr. Madhuri Gade, this vector encodes for an N-terminal hexa-histidine tag and confers ampicillin resistance. To insert the triple deletion in the scDH wildtype, gene assembly was performed employing the following primers: 5’-GCCGCGCGGCAGCCATATGGGATTATTTGC, 3’-GAGCTCGAATTCTTAAACGGGACGTCCTGC, 5’-TAAGAATTCGAGCTCCGTCGACA and 3’-CATATGGCTGCCGCGCGGCAC, resulting in the pET28a(+)-ΔscDH. The ΔMDH mutant was generated via PrimeStart® Max mutagenesis kit (TaKaRa Bio Inc), using primers 5’-GCGCTGGCGCTGCTGCTG and 3’-CAGCAGCGCCAGCGCGCCAATGCCCGCGCCCAGGTTCGCC, resulting in the pASK-IBA3(+)-ΔMDH construct.

*E. Coli* BL21 (DE3) cells (Merck) were transformed with the pET28a(+)-scDH and pET28a(+)-ΔscDH and grown in LB medium with 35 μg/ml kanamycin at 37 °C and shaken until an OD_600_ of ∼ 0.6. At this point, the protein expression was induced with 0.5 mM IPTG (final concentration) for a total of 5 h. The pASK-IBA3(+)-MDH construct was transformed in BL21 (DE3) cells and the pASK-IBA3(+)-ΔMDH in ArticExpress (DE3) cells, both cell lines were grown in LB medium with 100 35 μg/ml ampicillin at 37 °C and shaken until an OD_600_ of ∼ 0.6. Protein expression was induced with 200 ng/ml anhydrotetracycline for 16 h at 16 °C. Cells were harvested by centrifugation and stored at −80 °C. The pellets were resuspended in buffer A (20mM Tris pH 7.4, 150 mM NaCl and 25mM imidazole) supplemented with DNase, Lysozyme (Sigma) and EDTA-free protease inhibitors (Nakalai) and incubated on ice for 30 min. Cells were lysed by sonication at 30% amplitude for 1 minute (0.5 secs pulse, 0.5 secs pause). The resulting lysates were loaded onto a nickel affinity column HisTrap™ FF crude (5 ml) previously equilibrated with buffer A. After loading, the column was washed with 50 ml buffer A. An additional wash step was performed at 20% buffer B (buffer A plus 475 mM imidazole) and the target proteins eluted with an imidazole gradient. Fractions containing the target protein were collected and loaded onto a gel filtration column HiLoad 26/600 Superdex 75 pg (GE), previously equilibrated with size exclusion chromatography (SEC) buffer (20mM Tris pH 7.4 and 300 mM NaCl). All proteins eluted in a single peak.

### Crystallization and cryoprotection

The ΔscDH mutant was concentrated at 10 mg/ml in SEC buffer using a 30kDa molecular weight cutoff spin concentrator (Millipore). SAH was added at 1 mM to the concentrated protein sample. Crystallization screenings were performed by the sitting-drop vapor diffusion method. Drops were dispensed in 96-well plates using a mosquito high-throughput liquid handling robot (Mosquito® Xtal3). Each drop contained 400 nl protein solution plus 400 nl mother liqueur equilibrated against 50 ul mother liqueur reservoir. Crystals grew within a week from 6% v/v 2-propanol, 0.1 M sodium acetate trihydrate pH 4.5 and 26% v/v polyethylene glycol. The resulting crystals were cryoprotected in a 1:1 SEC buffer to mother liqueur solution, containing 1mM SAH and 20 % ethylene glycol prior to their flash-cooling in liquid nitrogen. The ΔMDH mutant was crystallized as the ΔscDH mutant described above. Crystals grew from 35% (v/v) MPD, 100 mM sodium cacodylate/hydrochloric acid (pH 6.5) and 50 mM zinc acetate within 2 weeks. The resulting crystals were cryoprotected in a 1:1 SEC buffer to mother liqueur solution, containing 1mM SAH and 20% ethylene glycol prior to their flash-cooling in liquid nitrogen.

### Data collection and processing

Complete data sets were collected on beamline BL32XU at Spring8 Radiation Institute, employing the automated data collection ZOO ^43^. Data processing was assessed with KAMO ^44^ automated processing from single crystals. The ΔscDH crystals belonged to space group 5 (C 1 2 1) with unit cell dimensions: a = 136.38, b = 82.76 and c = 92.66 Å. The unit cell contained one tetrameric copy in the asymmetric unit. The ΔMDH crystals belong to space group 19 (P2 21 21) with unit cell dimensions: a = 52.64 b = 98.35 and c = 118.56 Å. The asymmetric unit contained a protein dimer. Data collection statistics are given in **Table S2** and **S3**.

### Structure determination and refinement

The holo ΔscDH and apo ΔMDH structures were solved by molecular replacement with the Phaser crystallographic package, using the structure of TT0321 from *Thermus thermophilus* HB8 (PDB 2D1Y) and malate dehydrogenase from E. *coli* (PBB 1EMD) as search models, respectively. To avoid bias during refinement, the β1−loop−α1 regions were manually removed from the PDB models using PyMOL. The missing residues were built manually using Coot ^45^ during the refinement rounds with Phenix ^46^. The SAH cofactor was manually built into the refined structure to fit the electron density. Data processing and refinement statistic are given in in **Table S2** and **S3**.

### Isothermal titration calorimetry (ITC)

All titration experiments were performed at 25 °C using a Malvern MicroCal PEAQ-ITC microcalorimeter in ITC buffer (20mM Tris pH 8, 300 mM NaCl and 10% glycerol). Following concentrations of titrants and proteins were employed: 2 mM SAH to 65 μM scDH, 6 mM SAH to 160 μM ΔscDH, 1 mM SAH to 58 μM MDH, 500 μM SAH to 10 μM ΔMDH, 1 mM NAD to scDH, 6 mM NAD to 190 μM ΔscDH, 5 mM NAD to 10 μM MDH, and 5 mM NAD to ΔMDH. The protein concentration was calculated from the UV absorption at 280 nm with a Nanodrop spectrometer (Thermo Fisher Scientific) using the extinction coefficient obtained from the amino acid sequences using ExpaSy ProtParam tool ^47^ 52. The raw ITC data was analyzed with the ITC-Origin software using a “one-site” binding model after correction for the dilution heat of the compounds.

### Molecular dynamics

The MD simulations were performed using Gromacs version 2020.1 ^48^ and the charmm36-mar2019 force field ^49^. The two models used for the simulations were the short-chain dehydrogenase/reductase from *Thermus thermophilus* (scDH, PDB structure 2D1Y), and the crystal structure of the deletion mutant that was obtained in this work. To get the structure of the scDH in complex with SAH, the model was overlapped with the SAH-binding mutant and the position of the coenzymes was transferred. For each model, the system was solvated and neutralized with NaCl atoms in a dodecahedral box. The system was then equilibrated on energy, temperature, and pressure before performing five repeats of 200 ns of the production simulations. The simulations used the Verlet cut-off scheme for non-bonded interactions ^50^, the Particle Mesh Ewald for long-range electrostatic interactions ^51^, and the LINCS constraint algorithm ^52^. To analyze the trajectories, in-house python scripts together with Gromacs standard analysis package were used.

## Autor contributions

S.T.P. and PL conceived the project. S.T.P. designed the bioinformatical pipeline for sequence and structural analyses, expressed and purified proteins for structure solving. G.U. and S.T.P. conducted protein expression and their binding analysis with ITC. S.P. performed the molecular dynamics simulations. S.T.P., S.P., and P.L. analyzed the data. S.T.P. and P.L. wrote the manuscript with inputs from S.P. This project was supervised by PL.

## Acknowledgements

This work has been supported by OIST funding. We thank Andrei Lupas, Liam Longo and Benjamin Clifton for insightful comments on this manuscript. We are grateful for the help provided by the Scientific Computing, the Data Analysis and the Instrumental Analysis sections at OIST. We also thank the Japan Synchrotron Radiation Research Institute for their assistance with automated data collection, in particular to Naoki Sakai for his excellent technical support.

