## Supplementary material for "Insertions and deletions mediated functional divergence of Rossmann fold enzymes": Suplemental Information: 20220516_SI.pdf

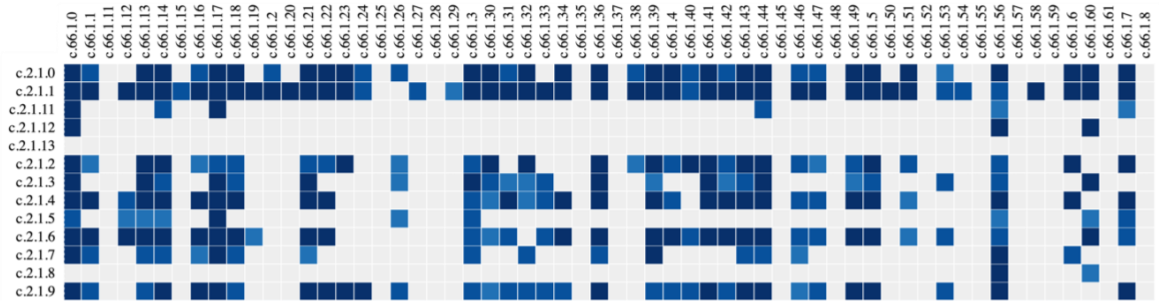

**Figure S1. Homology connections between Rossman: NAD(P)-binding oxidoreductase and SAM-dependent methyltransferase superfamilies.** The present matrix illustrates the distant homology connections between protein families belonging to SCOP superfamilies: “NAD(P)-binding Rossmann fold domains” (SCOP ID: c.2) and “S-adenosyl-L-methionine-dependent methyltransferases” (SCOP ID: c.66). Squares in gray represent an HHsuite probability below 80%. Blue squares represent connections that show HHsuite probabilities ranging from 80% (light blue), over 90% (middle-tone blue) up to 100% (dark blue). This matrix has been generated, using the Fuzzle webserver [1].

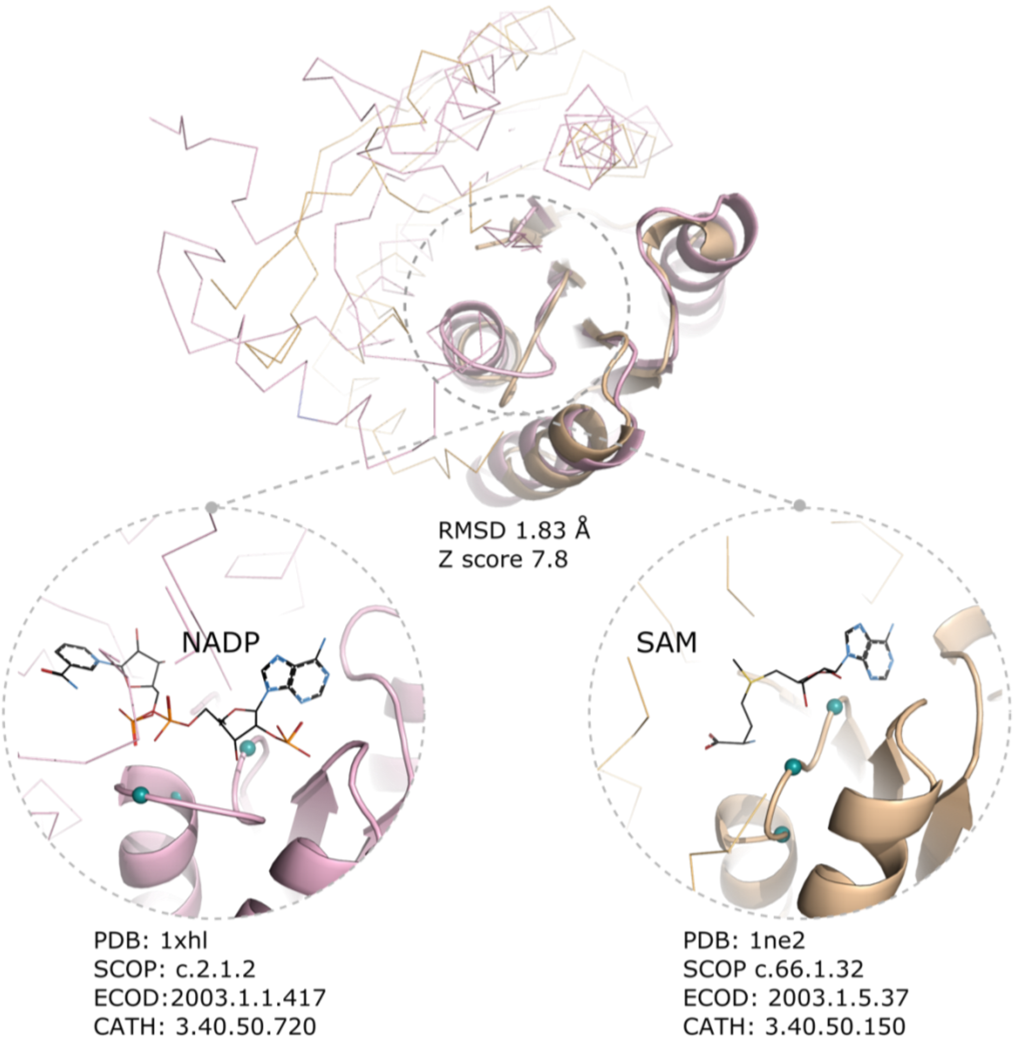

```
>d1xhla_ c.2.1.2 (A:) Putative Tropinone Reductase II {Nematode (Caenorhabditis elegans) [TaxId: 6239]}
Probab=90.98  E-value=0.033  Score=47.81  Aligned_cols=67  Identities=40%  Similarity=0.484  Sum_probs=0.0

Q ss_pred               CCCEEEEeCCCCCH---HHHHHHHcCCCEEEEEEECCHHHHHHHHHHc-------CCCceEEccccccC-----------
Q d1ne2a_          47 IGGRSVIDAGTGNGI---LACGSYLLGAESVTAFDIDPDAIETAKRNC-------GGVNFMVADVSEIS-----------  105 (197)
Q Consensus        47 i~gk~VlDlGcGtG~---l~i~a~~~ga~~V~gvDiD~~al~~A~~n~--------~~i~~i~~Di~~~-----------  105 (197)
                        ++||++|==|.++|+   .+...+..||+ |+.+|.|++.++.+.+.+       .++.++.+|+++..
T Consensus         2 l~gKvalVTGas~GIG~aiA~~fa~eGA~-Vvi~~r~~~~l~~~~~ei~~~g~~~~~~~~~~~~~~~Dv~~~~~~v~~~v~~~~~~   80 (274)
T d1xhla_           2 FSGKSVIITGSSNGIGRSAAVIFAKEGAQ-VTITGRNEDRLEETKQQILKAGVPAEKINAVVADVTEASGQDDIINTTLA   80 (274)
T ss_pred               CCCEEEEcCCCCHHHHHHHHHHHHHCCCE-EEEEECCHHHHHHHHHHHHHcCCChhceEEEECCCCCHHHHHHHHHHHHH

Q ss_pred               -CCccEEEEc
Q d1ne2a_         106 --GKYDTWIMN  114 (197)
Q Consensus       106 --~~~D~Vi~N  114 (197)
                         ++.|+.|-|
T Consensus        81 ~fG~iDiLVNN   91 (274)
T d1xhla_          81 KFGKIDILVNN   91 (274)
T ss_pred               HCCCCEEEeC
```

**Figure S2. An example of local homology between NAD- and SAM-dependent Rossmann domains.** Distant homology can be detected between Rossmann: NAD(P)-dependent oxidoreductases and SAM-binding methyltransferases[2]. Here, we display the sequence-based profile alignment, corresponding to the top-scored pair obtained from the HHsuite searches taking the sequence identity as the main criteria. The structure of a putative tropinone reductase (pink), which has been crystallized with NADP and tropinone, shares 40% sequence identity along 67 residues with a hypothetical methyltransferase (yellow). This similarity corresponds also in structure (RMSD of 1.2 Å and a Z score of 7.8). Sequence similarity was predicted using HHsuite [3] and structural alignments were assessed with PDBeFold [4]. SAM has been superimposed from PDB 1NV8. Glycine-rich motives are shown in turquoise. Abbreviations: Probab = HHsuite probability, Q = query, T = target, C = coil/loop, E = beta strand, H = helix.

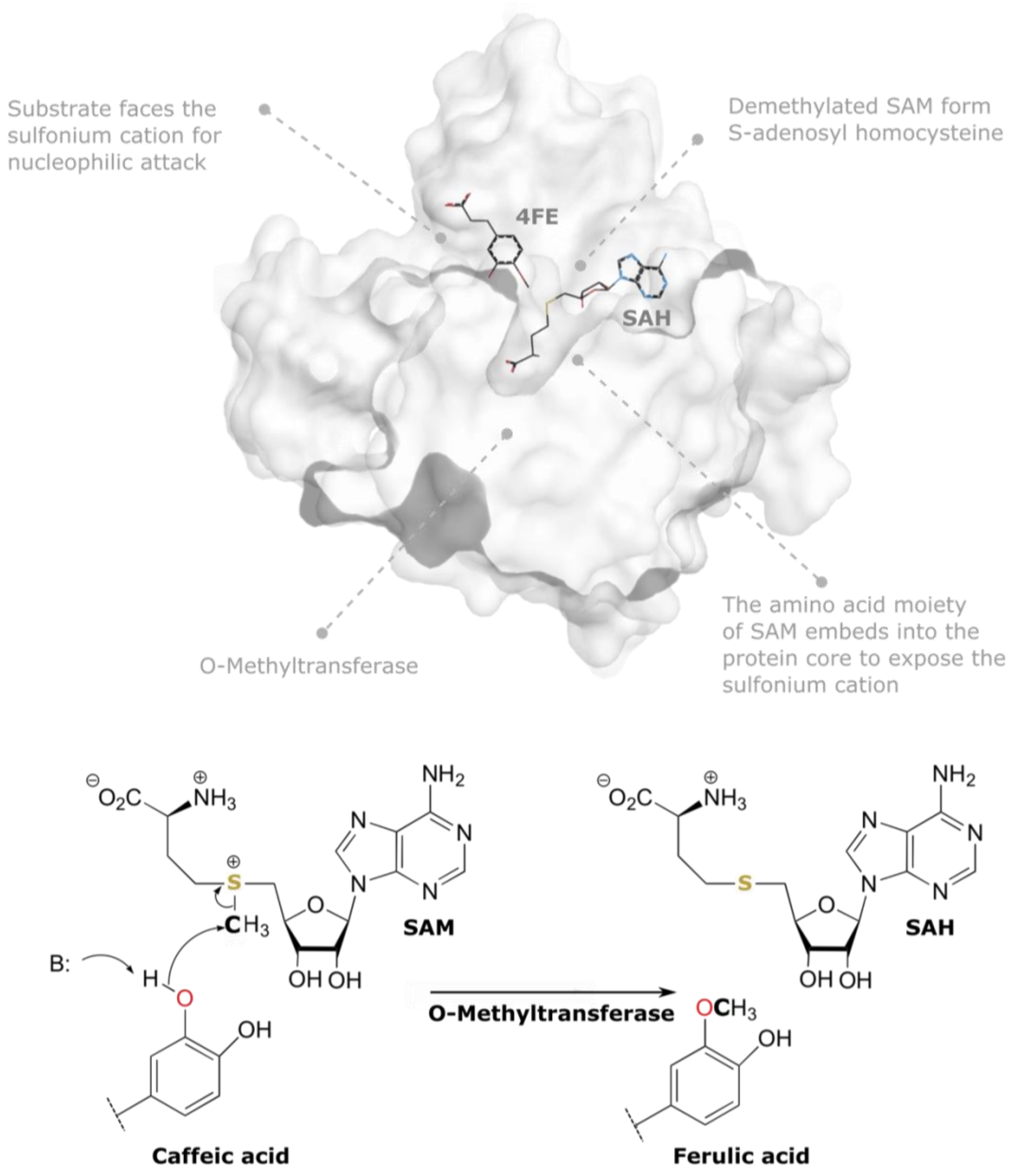

**Figure S3. Catalytically productive conformation of SAM.** In Rossmann methyltransferases, the coenzyme S-adenosyl methionine adopts a bended conformation, where its amino acid moiety is embedded inside the enzyme core. This conformation facilitates the exposure of the sulfonium cation towards the substrate. As an example, here we illustrate the reaction mechanism of an O-methyltransferase from *Cyanobacterium Synechocystis* (PDB 3CBG), which has been crystallized with the demethylated coenzyme and (SAH) and its methylated substrate ferulic acid. The reaction mechanism initiates by the extraction of a proton from the reacting hydroxy group, followed by a nucleophilic attack towards the SAM methyl group, in an SN2 reaction mechanism.

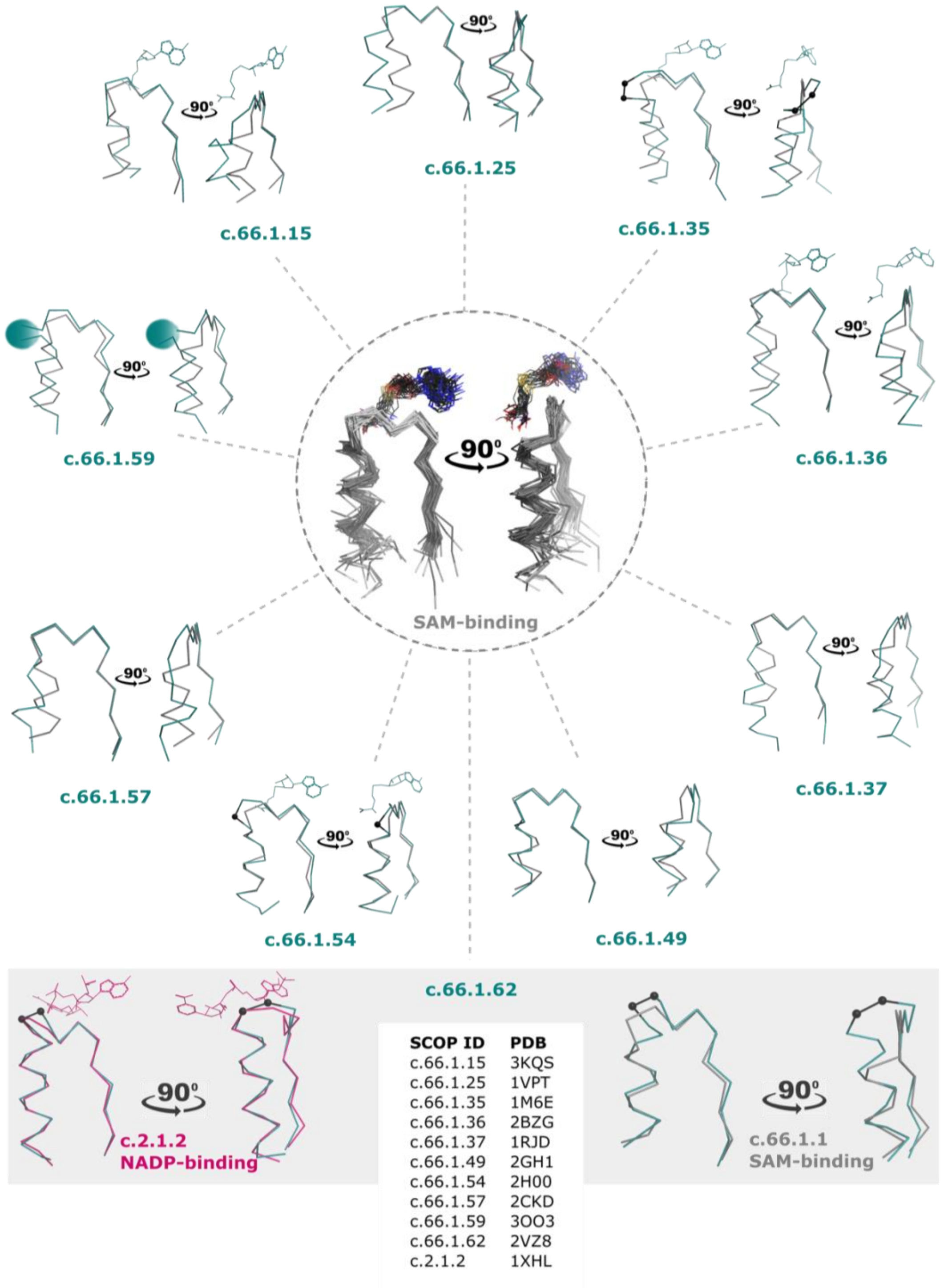

| SCOP ID | PDB |
|---|---|
| c.66.1.15 | 3KQS |
| c.66.1.25 | 1VPT |
| c.66.1.35 | 1M6E |
| c.66.1.36 | 2BZG |
| c.66.1.37 | 1RJD |
| c.66.1.49 | 2GH1 |
| c.66.1.54 | 2H00 |
| c.66.1.57 | 2CKD |
| c.66.1.59 | 3OO3 |
| c.66.1.62 | 2VZ8 |
| c.2.1.2 | 1XHL |

**Figure S4. The highly conserved β1–loop– α1 structure of the SAM coenzyme-binding pocket of dependent methyltransferases against its outliers.** 49 out of 60 SAM-dependent methyltransferase SCOP families share similar β1–loop–α1 structural features at the coenzyme-binding pocket (center). Outlier families are displayed surrounding the main motif. Some of the structural outliers present insertions (black spheres). Family c.66.1.59 presents a small domain insertion (represented as a turquoise sphere). Family c.66.1.62 (bottom), displays similar β1–loop–α1 structural features as the NAD(P) coenzyme-binding pocket. This family has been reported as non-catalytically active but sorted within the SAM-dependent methylase superfamily by SCOP. SCOP PDB IDs used for this figure are listed as a table.

a)

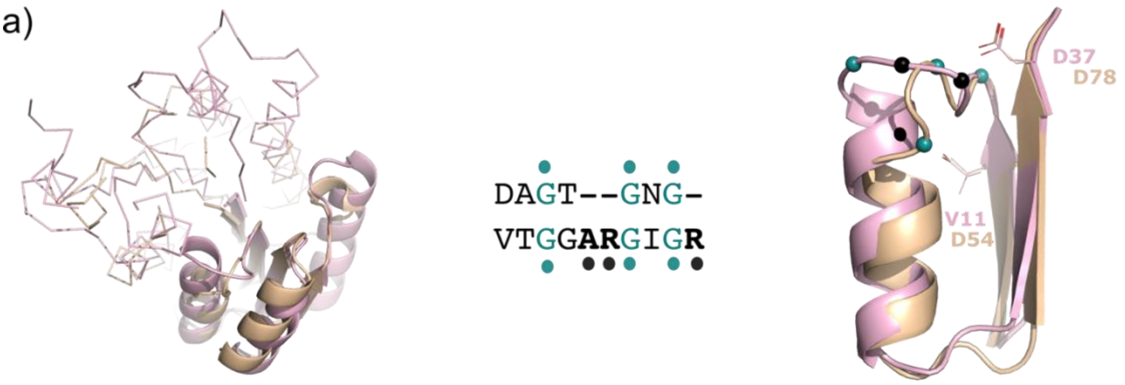

```
Query SCOP ID: d2d1ya1 c.2.1.2 (A:2-249) Oxidoreductase TT0321 from Thermus thermophilus
Probab=91.56  E-value=0.025  Score=48.01  Aligned_cols=67  Identities=30%  Similarity=0.451  Sum_probs=0.0

Q ss_pred              CCCCCEEEEeCCCCCH---HHHHHHHcCCCEEEEEEECCHHHHHHHHHHcCCCceEEccccccC-------------CCc
Q d1ne2a_           45 GNIGGRSVIDAGTGNGI---LACGSYLLGAESVTAFDIDPDAIETAKRNCGGVNFMVADVSEIS-------------GKY  108 (197)
Q Consensus         45 gdi~gk~VlDlGcGtG~---l~i~a~~~ga~~V~gvDiD~~al~~A~~n~~~i~~i~~Di~~~~-------------~~~  108 (197)
                         |-++||+++==|.++|+    .+...+..||+ |+..|++++..+.+++.  +..++.+|+++..              ++.
T Consensus          1 Grl~gKvaiVTGa~~GIG~aia~~fa~eGA~-Vvi~d~~~e~~~~~~~~~~~~~~~~~DVt~~~~v~~~v~~~~~~~~G~i   77 (248)
T d2d1ya1           1 GLFAGKGVLVTGGARGIGRAIAQAFAREGAL-VALCDLRPEGKEVAEAI--GGAFFQVDLEDERERVRFVEEAAYALGRV   77 (248)
T ss_pred              CCCCCEEEECCCCCHHHHHHHHHHHHCCCE-EEEEECCHHHHHHHHHh--CCEEEECCCCHHHHHHHHHHHHHHHHCCC

Q ss_pred              EEEEc
Q d1ne2a_          109 DTWIMN  114 (197)
Q Consensus        109 D~Vi~N  114 (197)
                         |+.|-|
T Consensus         78 DiLvNN   83 (248)
T d2d1ya1          78 DVLVNN   83 (248)
T ss_pred              EEEEC
```

b)

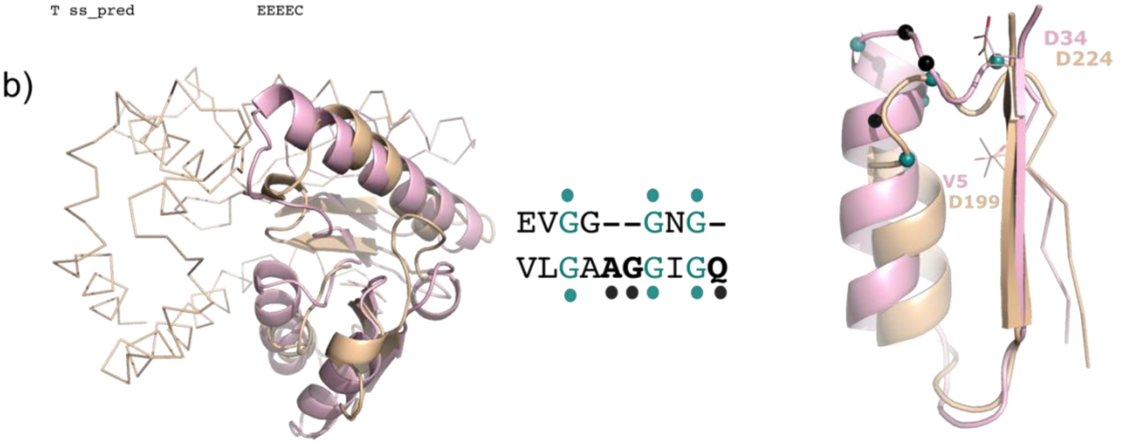

```
Query SCOP ID: d3hhpa1 c.2.1.0 (A:1-145) malate dehydrogenase from Escherichia coli K-12 [TaxId: 83333]}
Probab=52.97  E-value=4.1  Score=31.06  Aligned_cols=93  Identities=19%  Similarity=0.280  Sum_probs=0.0

Q ss_pred              c--CCcHHHHHHHHH-HCCCc-EEEEccCCceee-------eccCCcccc-eecccccccCccchhhhhhHhhcCCH-
Q d2qyoa2           87 VG--GGNGTTGKIICE-TFPKL-TCVVFDRPKVVE-------NLCGSNNLT-YVGGDMFISVPKADAVLLKAVLHDWTD-  153 (244)
Q Consensus         87 vG--GG~G~~~~~i~~~~P~l~~iv~Dlp~vi~~~~~~~~~~a~~~~rv~~~~~gDff~~~P~aDvy~l~~ilHdw~d-  153 (244)
                         ||   |..|...+.+++ +.|.. ..+++|..+..+          ++....++. +...|-++.+..+|++++..=...=+.
T Consensus          6 iGaaG~vG~~~a~~L~~~~~~~~~~el~L~D~~~~~~g~a~Dl~~~~~~~~~~~~~~~~~~~~~~~al~~aDiVvitag~~~~~~~~   85 (145)
T d3hhpa1           6 LGAAGGIGQALALLLKTQLPSGSELSLYDIAPVTPGVAVDLSHIPTAVKIKGFSGEDATPALEGADVVLISAGVARKPGM   85 (145)
T ss_pred              CCCChHHHHHHHHHHccCCCCCEEEEEeCCCCChhheeecccCCCccccceEEecCCchHHHCCCcEEEEecCcCCCCCC

Q ss_pred              -----HHHHHHHHHHHhCCCCCCCceEEEE
Q d2qyoa2          154 ------KDCIKILKKCKEAVTSDGKRGKVIVI  179 (244)
Q Consensus        154 ------~~~~~IL~~~~~Al~~~~~~grlII~  179 (244)
                               +.+..|++.+.+++....+.+.++|+
T Consensus         86 sR~dl~~~N~~iv~~i~~~~i~~~~~p~aivivv  117 (145)
T d3hhpa1          86 DRSDLFNVNAGIVKNLVQQVAKTCPKACIGII  117 (145)
T ss_pred              hhhHHHHHHHHHHHHHHHHHHHHCCCceEEEe
```

**Figure S5. Structural and sequence alignments of selected targets for design.** Structural alignments for oxidoreductases (pink): PDB 2D1Y (a) and 3HHP (b) against their best-ranked SAM-binder hits (yellow). In carton, are shown the homology regions detected by HHsuite and in ribbon, regions where homology cannot be detected at a sequence level. Glycine motifs are colored in turquoise and deleted regions are highlighted as black spheres. HHsuite alignments are displayed below the structures. Abbreviations: Probab = HHsuite probability, Q = query, T = target, C = coil/loop, E = beta strand, H = helix.

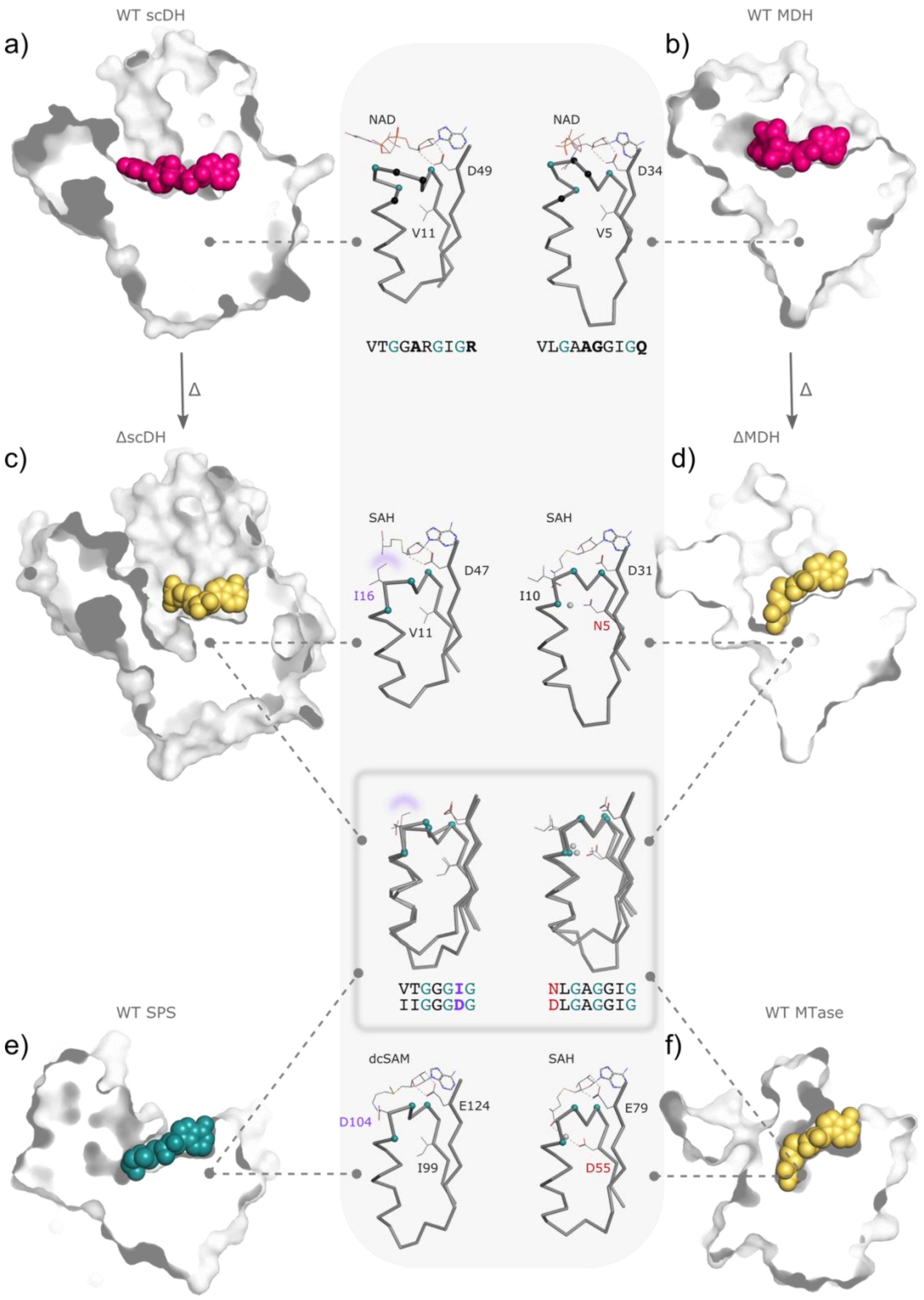

**Figure S6. Structural comparison of the engineered proteins against natural NAD(P)-, SAM- and dcSAM-binding enzymes.** The crystal structures (surface) of **(a)** WT short chain dehydrogenase/reductase (scDH, PDB 2D1Y) and **(b)** WT malate dehydrogenase (MDH, PDB 1EMD) in complex with NAD (pink spheres) show the horizontal shape of the coenzyme-binding pocket. Their $\beta1-\alpha1-\beta2$ structural elements (ribbon) display the bidentate interaction of Asp49/34 to the ribosyl moiety and a hydrophobic residue (Val11/Val5), typically found at position one of the NAD(P)-binding motif. InDels (black) and Gly-rich motifs (turquoise) are shown as spheres. (**c**) Deletion of three amino acids (black spheres) from scDH resulted in the mutant ΔscDH (PDB 7XQM), whose crystal structure (surface) in complex with demethylated SAM (SAH) revealed a larger pocket than that of the WT scDH, however obstructed by residue Ile16 (purple) that impedes binding to SAM in its catalytically productive bent conformation. The bidentate interaction of the ribosyl moiety of SAH to Asp47 and the hydrophobic residue Val11 are shown as lines. **(d)** Deletion of three amino acids (black spheres) and a single substitution V5N (red) on the WT MDH resulted in mutant ΔMDH (PDB 7XQN), whose crystal structure (surface) revealed the typical shape of the SAM-binding pocket, providing enough space for its catalytically productive conformation. The demethylated SAM coenzyme (SAH, yellow spheres) was superimposed from PDB 1WY7. The ΔMDH β1–α1–β2 structural elements show the bidentate interaction of the ribosyl moiety to Asp31 and the hydrogen-bonding interaction of Asn5 with a water molecule, as observed for other WT SAM-binding enzymes. In contrast to Ile16 in ΔscDH, Ile10 (lines) in ΔMDH does not obstruct the coenzyme pocket, displaying an inverted side chain conformation. (**e**) WT spermidine synthase (SPS, PDB 2O0L) bound to decarboxylated SAM (dcSAM, turquoise spheres) shows a horizontal coenzyme pocket similar to that of NAD. The SPS β1–α1–β2 structural elements display a hydrophobic residue (Ile99) at position one of the coenzyme-binding motif in analogy to Val11 in the scDH, and WT oxidoreductases, instead of the charge residues (Asp/Glu) mainly observed for other SAM-binding enzymes. Similar to Ile16 in ΔscDH, Asp104 (purple) blocks the potential binding to SAM in its bent conformation. Superposition of the β1–α1–β2 structural elements of SPS and ΔscDH (gray square) illustrate their resemblance also at a structural level. (**f**) The crystal structure (surface) of a putative methyltransferase (MTase, PDB 1WY7) in complex with SAH (yellow spheres) illustrates the canonical shape of the SAM-binding pocket that allows its bent conformation. The MTase MDH β1–α1–β2 structural elements bound to SAH (lines) shows the bidentate interaction of its ribosyl moiety to Glu77 (lines) and the charged residue D55 (red) interacting via hydrogen binding with a water molecule. Superposition of the β1–α1–β2 structural elements of MTase and ΔMDH (gray square) illustrates almost identical structural features, including: the hydrogen bonding of Asp55/Asn5 to a water molecule and the same orientation of their Gly-rich motifs (turquoise spheres).

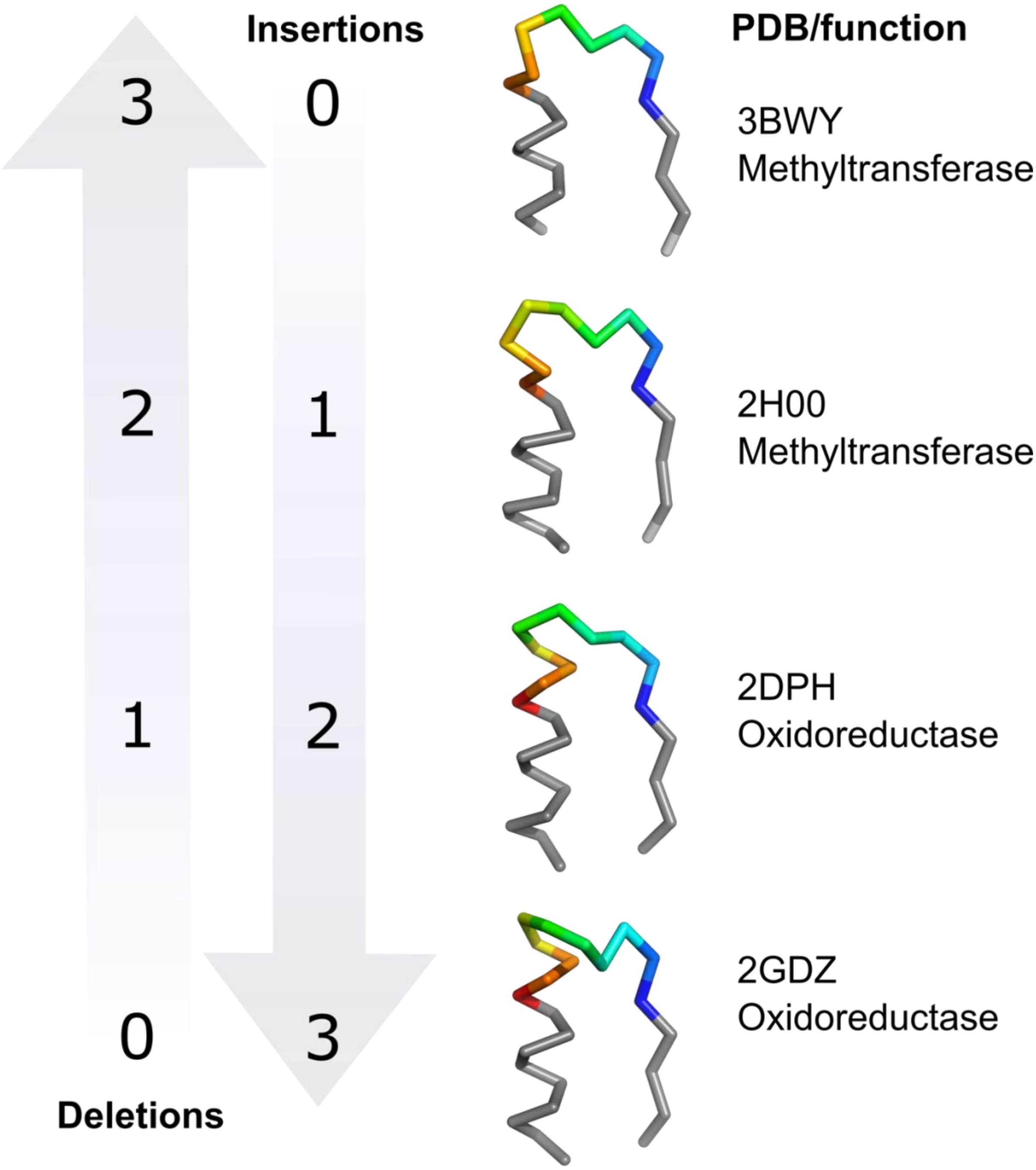

**Figure S7. InDel intermediates.** Parting from the typical β1–loop-α1 structural features of methyltransferases, the catechol O-methyltransferase (PDB 3BWY) exemplifies the typical shorter helix (0 insertions) of methyltransferases (top) compared to oxidoreductases (PDB 2GDZ, bottom). Intermediate β1–loop-α1 structures can be found for both, methyl transferases and oxidoreductases. As an example, a human methyltransferase (PDB 2H00) presents one amino acid insertion at the coenzyme-binding loop. Similarly, an NAD-binding formaldehyde dismutase (PDB 2DPH), presents a deletion of one amino acid and a shorter helix α1 compared to the most abundadnt β1–loop–α1 structure of oxidoreductases (**Figure 1e**). This biding motif is adopted by 32% of the domains belonging to the NAD(P)-binding superfamily (according to SCOP release 2.08 at 40% sequence identify) and is related in sequence and structure to the β1–loop–α1 coenzyme-binding motif of Rossmann flavin adenine dinucleotide (FAD)-binding domains.

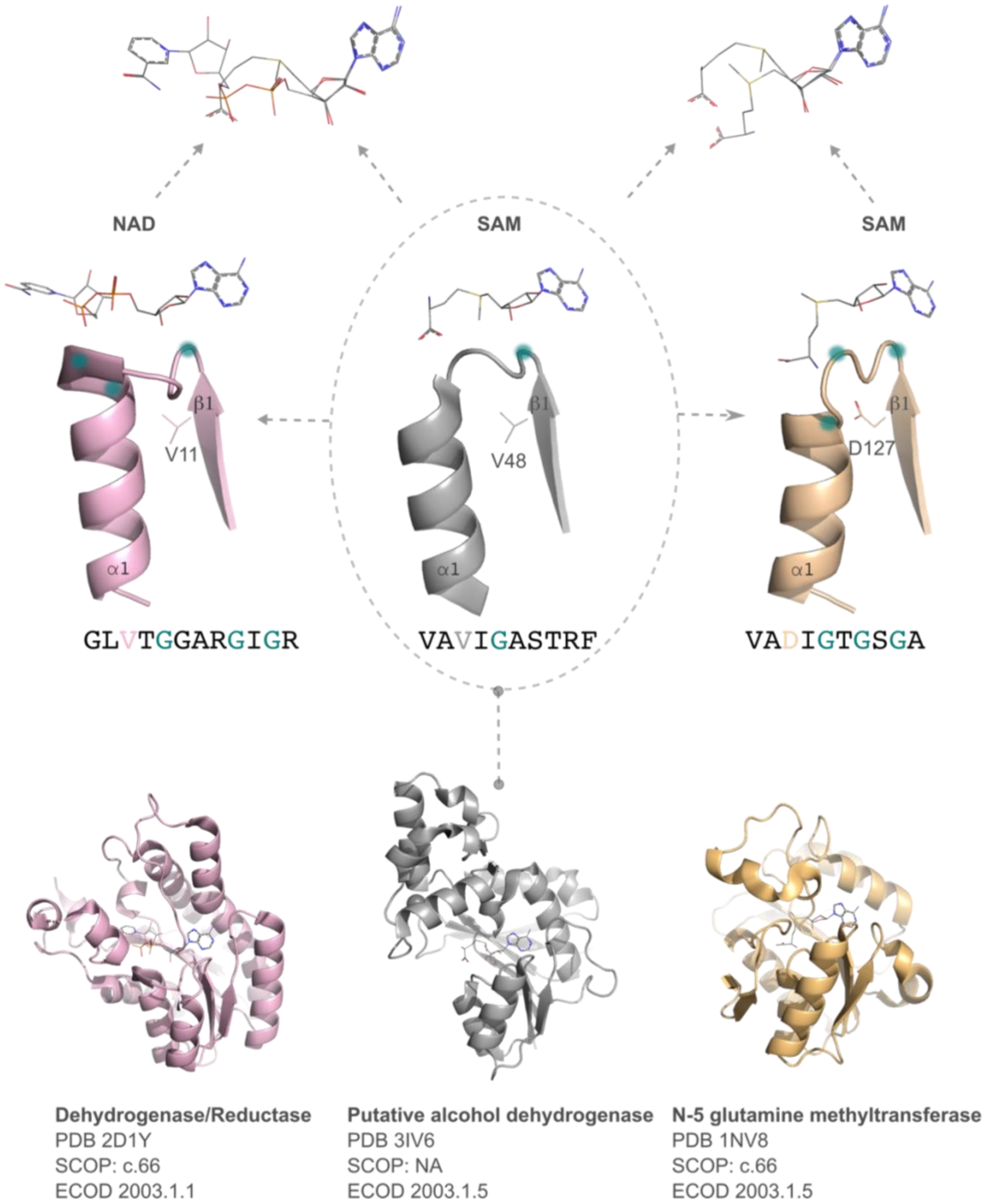

**Figure S8 An example of a SAM/NAD structural intermediate**. A putative Zn-dependent alcohol dehydrogenase (gray) crystallized with SAM presents $\beta1$–loop–$\alpha1$ intermediate features (dashed oval) compared to oxidoreductases (pink) and methyltransferases (yellow). For instance, the beta strand $\beta1$ lacks the charged residue that stabilized SAM in its catalytically productive bent conformation in methyltransferases. As a result, the sulfonium cation points downwards, being not available for methylation. In addition, the helix $\alpha1$ displays an intermediate length and the coenzyme-binding loop a distinct shape compared to both, NAD, and SAM-binding superfamilies. This protein is classified by ECOD as methyltransferase. However, it has not been functionally characterized. NAD and SAM-binding Gly-rich motifs are displayed as turquoise circles.

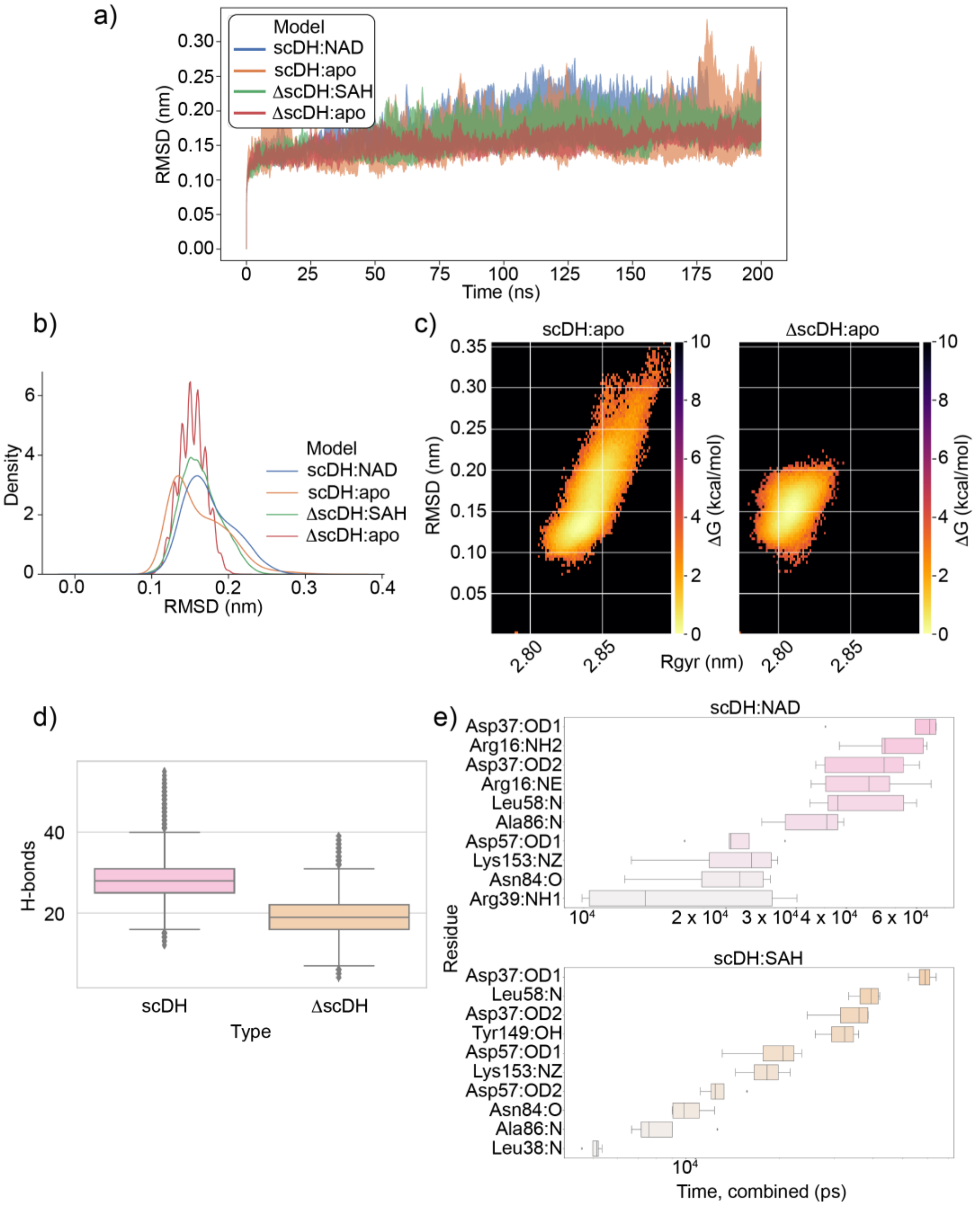

**Figure S9. Additional Molecular Dynamics analysis of coenzyme binding for the short chain dehydrogenase and the deletion mutant.** (**a**) The average Root Mean Square Deviation (RMSD) in a window of 40 residues is plotted every fifth time step. The line represents the average of the four repeats, while the width represents the standard deviation. (**b**) The distribution of RMSD per simulation type. All four simulations show a stable, single, main conformation as indicated by the single peak. (**c**) The Free Energy Surface (FES) of the two apo simulations, obtained by combining the five repeats and extracting the RMSD and Radius of Gyration (Rgyr) for each time step of the trajectory. The WT scDH appears to sample a higher number of conformations compared to the ΔscDH mutant. (**d**) The total number of hydrogen bonds between protein and coenzyme per time step of the two holo simulation types. The scDH simulations with four NAD coenzymes show on average a higher number of H-bonds than the ΔscDH simulations with four SAH. (**e**) The total time spent in a hydrogen bond for each residue/atom pair. The two simulation types – scDH with four NAD and scDH with four SAH – were compared. Arg16 does not seem to play an important role in SAH binding.

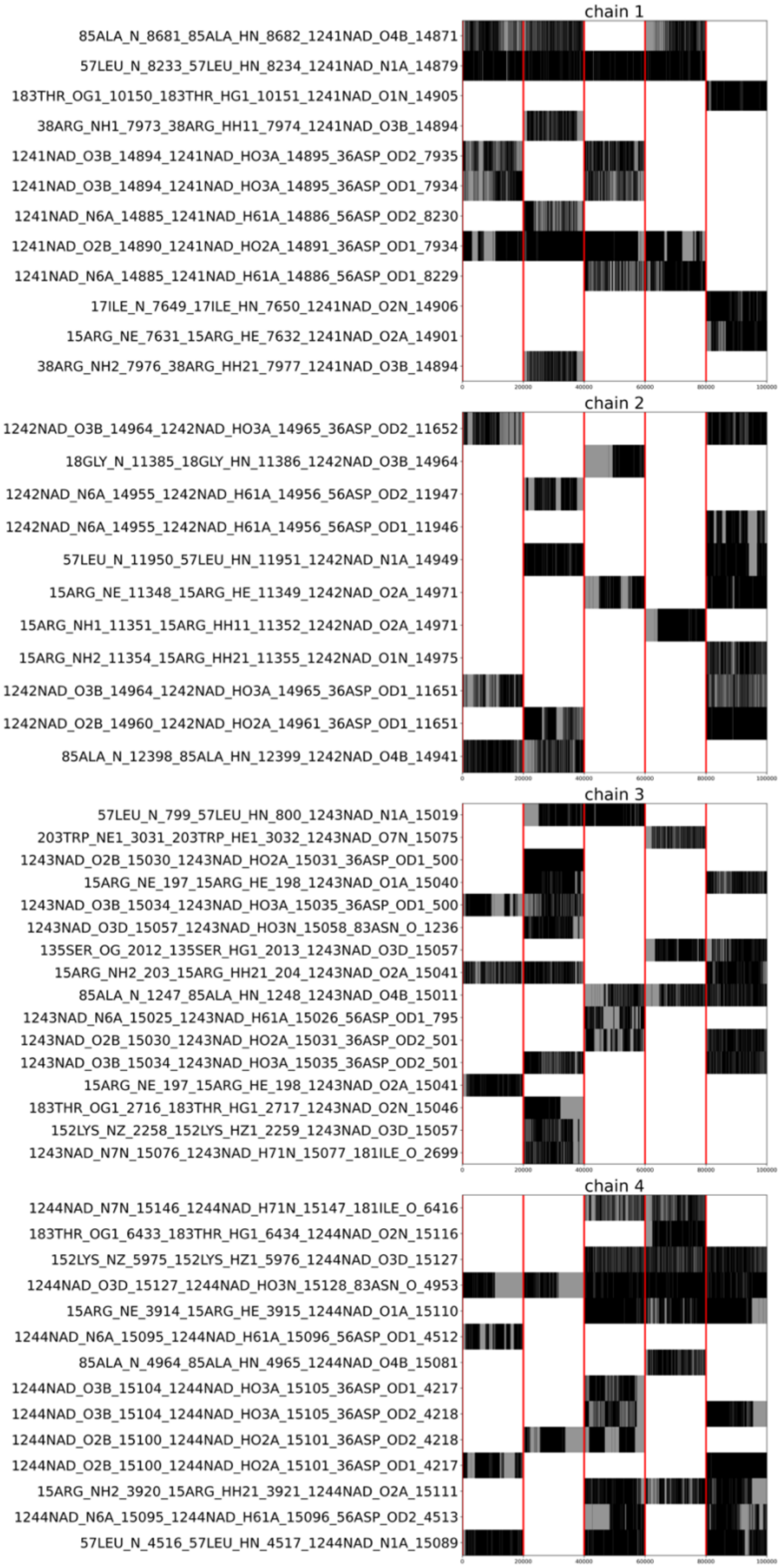

**Figure S10. Hydrogen bond permanence per simulation repeat of the WT scDH.** The Hydrogen bonds that were found for at least 50% of the simulation time in one repeat are reported. On the X axis, the simulation time is divided in the five repeats using red lines. The black lines show when the hydrogen bond is present. The results for each of the four chains are shown.

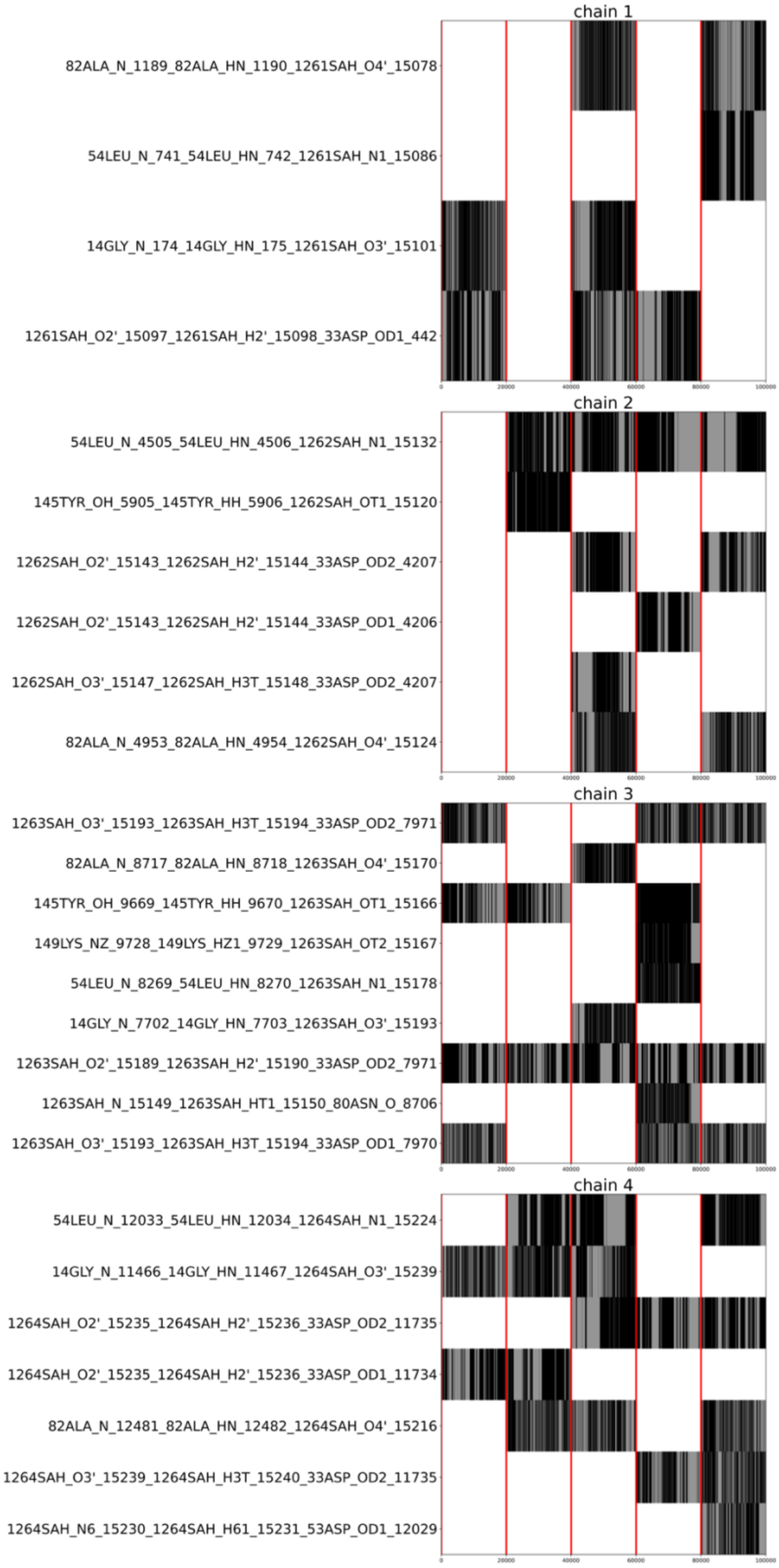

**Figure S11. Hydrogen bond permanence per simulation repeat of the ΔscDH mutant.** The Hydrogen bonds that were found for at least 50% of the simulation time in one repeat are reported. On the X axis, the simulation time is divided in the five repeats using red lines. The black lines show when the hydrogen bond is present. The results for each of the four chains are shown.

**Table S1. Ranking of best candidates for InDel engineering**. Top-ranked targets were obtained through sequence-base profile comparisons with HHsuite [3], employing the astral compendium of SCOP database 2.07 (March 2019) release (please refer to methods section for more details). Methyltransferase domains were used as queries to obtain their corresponding oxidoreductase hits. The obtained alignments were ranked according to sequence identity (30% o and up) and filtered according to their length (60 residues and up). Targets used for experimental characterization were selected due to thermostability or high expression yields (highlighted in gray). The additional hits showed below the table resulted from manual inspection of well characterized enzymes.

| Rank | PDB query | SCOP ID | PDB hit | SCOP ID | Sequence identity(%) | aligned residues |
|---|---|---|---|---|---|---|
| 1 | 1ne2 | d1ne2a_ | 1xhl | d1xhla_ | 40 | 63 |
| 2 | 1ne2 | d1ne2a_ | 1spx | d1spxa_ | 34 | 71 |
| 3 | 1ne2 | d1ne2a_ | 1iy8 | d1iy8a_ | 31 | 64 |
| 4 | 1nv8 | d1nv8a_ | 4i5f | d4i5fa_ | 31 | 67 |
| 5 | 1ne2 | d1ne2a_ | 2d1y | d2d1ya1 | 30 | 67 |
| 6 | 1uir | d1uira_ | 1jvb | d1jvba2 | 30 | 71 |
| 7 | 1wy7 | d1wy7a1 | 2gdz | d2gdza1 | 30 | 71 |
| 8 | 1wy7 | d1wy7a1 | 3o38 | d3o38a_ | 30 | 64 |
| 9 | 1wy7 | d1wy7a1 | 3n74 | d3n74a_ | 30 | 73 |
| 10 | 2b2c | d2b2ca1 | 2a35 | d2a35a1 | 30 | 76 |
| 11 | 1f3l | d1f3la_ | 3n74 | d3n74a_ | 30 | 67 |
| 12 | 2b3t | d2b3ta_ | 3t7c | d3t7ca_ | 30 | 67 |
| 13 | 2b3t | d2b3ta_ | 3uwr | d3uwra_ | 30 | 69 |
| 14 | 4c08 | d4c08a_ | 4j2h | d4j2ha_ | 30 | 70 |
| 15 | 4c08 | d4c08a_ | 3awd | d3awda_ | 30 | 67 |
| 16 | 4hc4 | d4hc4a_ | 4j2h | d4j2ha_ | 30 | 94 |
| 17 | 4hc4 | d4hc4a_ | 3o38 | d3o38a_ | 30 | 67 |
| | 2d1y | d2d1ya1 | 2nxc | d2nxca1 | 32 | 66 |
| | 2qyo | d2qyoa2 | 3hhp | d3hhpa1 | 19 | 93 |

**Table S2 Data collection and refinement statistics for ΔscDH in complex with SAH**

| | |
|---|---|
| WAVELENGTH | 1.000 Å |
| RESOLUTION RANGE | 44.01 - 2.71 (2.807 - 2.71) |
| SPACE GROUP | C 1 2 1 |
| UNIT CELL | 136.38 82.76 92.66 90 111.995 90 |
| TOTAL REFLECTIONS | 50905 (4339) |
| UNIQUE REFLECTIONS | 25801 (2271) |
| MULTIPLICITY | 2.0 (1.9) |
| COMPLETENESS (%) | 98.7 (87.35) |
| MEAN I/SIGMA(I) | 21.4 (11.44) |
| WILSON B-FACTOR | 26.42 |
| R-MERGE | 0.02485 (0.04784) |
| R-MEAS | 0.03514 (0.06766) |
| R-PIM | 0.02485 (0.04784) |
| CC1/2 | 0.997 (0.99) |
| CC* | 0.999 (0.998) |
| REFLECTIONS USED IN REFINEMENT | 25791 (2271) |
| REFLECTIONS USED FOR R-FREE | 1999 (176) |
| R-WORK | 0.18 (0.20) |
| R-FREE | 0.18 (0.2948) |
| CC(WORK) | 0.944 (0.894) |
| CC(FREE) | 0.940 (0.901) |
| NUMBER OF NON-HYDROGEN ATOMS | 7660 |
| MACROMOLECULES | 7536 |
| LIGANDS | 0 |
| SOLVENT | 124 |
| PROTEIN RESIDUES | 1014 |
| RMS(BONDS) | 0.011 |
| RMS(ANGLES) | 1.53 |
| RAMACHANDRAN FAVORED (%) | 96.61 |
| RAMACHANDRAN ALLOWED (%) | 3.39 |
| RAMACHANDRAN OUTLIERS (%) | 0.0 |
| ROTAMER OUTLIERS (%) | 0.97 |
| CLASHSCORE | 1.71 |
| AVERAGE B-FACTOR | 22.92 |
| MACROMOLECULES | 22.92 |
| SOLVENT | 23.26 |

**Table S3 Data collection and refinement statistics for the ΔMDH apo structure.**

| | |
|---|---|
| WAVELENGTH | 1.000 Å |
| RESOLUTION RANGE | 49.17 - 1.98 (2.051 - 1.98) |
| SPACE GROUP | P 2 21 21 |
| UNIT CELL | 52.64 98.35 118.56 90 90 90 |
| TOTAL REFLECTIONS | 87402 (8626) |
| UNIQUE REFLECTIONS | 43701 (4313) |
| MULTIPLICITY | 2.0 (2.0) |
| COMPLETENESS (%) | 99.97 (99.93) |
| MEAN I/SIGMA(I) | 8.64 (1.55) |
| WILSON B-FACTOR | 22.09 |
| R-MERGE | 0.03141 (0.1771) |
| R-MEAS | 0.04442 (0.2505) |
| R-PIM | 0.03141 (0.1771) |
| CC1/2 | 0.998 (0.867) |
| CC* | 1 (0.964) |
| REFLECTIONS USED IN REFINEMENT | 43694 (4312) |
| REFLECTIONS USED FOR R-FREE | 2000 (197) |
| R-WORK | 0.1775 (0.2597) |
| R-FREE | 0.1864 (0.2629) |
| CC(WORK) | 0.960 (0.860) |
| CC(FREE) | 0.954 (0.818) |
| NUMBER OF NON-HYDROGEN ATOMS | 5007 |
| MACROMOLECULES | 4522 |
| LIGANDS | 0 |
| SOLVENT | 485 |
| PROTEIN RESIDUES | 620 |
| RMS(BONDS) | 0.010 |
| RMS(ANGLES) | 1.29 |
| RAMACHANDRAN FAVORED (%) | 99.03 |
| RAMACHANDRAN ALLOWED (%) | 0.97 |
| RAMACHANDRAN OUTLIERS (%) | 0.0 |
| ROTAMER OUTLIERS (%) | 2.7 |
| CLASHSCORE | 1.41 |
| AVERAGE B-FACTOR | 23.59 |
| MACROMOLECULES | 22.79 |
| SOLVENT | 31.06 |

**Table S4 List of protein domains used for structural super positions.** 312 Structural representatives, corresponding to the SAM and (SAM-related)-binding domains, according to CATH database (Superfamily ID 3.40.50.150).

| No. | PDB | CATH ID | No. | PDB | CATH ID | No. | PDB | CATH ID |
|---|---|---|---|---|---|---|---|---|
| 1 | 1af7 | 1af7A02_A | 41 | 1ws6 | 1ws6A00_A | 81 | 2jjq | 2jjqA02_A |
| 2 | 1boo | 1booA00_A | 42 | 1wzn | 1wznA01_A | 82 | 2k4m | 2k4mA00_A |
| 3 | 1dl5 | 1dl5A01_A | 43 | 1x19 | 1x19A02_A | 83 | 2nxc | 2nxcA03_A |
| 4 | 1dus | 1dusA00_A | 44 | 1xxl | 1xxlA00_A | 84 | 2o57 | 2o57A02_A |
| 5 | 1eg2 | 1eg2A00_A | 45 | 1y8c | 1y8cA01_A | 85 | 2o57 | 2o57D00_D |
| 6 | 1ej0 | 1ej0A00_A | 46 | 1yb2 | 1yb2A00_A | 86 | 2ob1 | 2ob1A00_A |
| 7 | 1ej6 | 1ej6A04_A | 47 | 1yf3 | 1yf3A01_A | 87 | 2okc | 2okcA02_A |
| 8 | 1f3l | 1f3lA01_A | 48 | 1yf3 | 1yf3A01_C | 88 | 2ozv | 2ozvA01_A |
| 9 | 1fp1 | 1fp1D02_D | 49 | 1yf3 | 1yf3A01_D | 89 | 2p35 | 2p35A01_A |
| 10 | 1fp2 | 1fp2A02_A | 50 | 1yzh | 1yzhB00_B | 90 | 2p7i | 2p7iA00_A |
| 11 | 1g55 | 1g55A02_A | 51 | 1zkd | 1zkdA02_A | 91 | 2p8j | 2p8jA00_A |
| 12 | 1g60 | 1g60B00_B | 52 | 2a14 | 2a14A00_A | 92 | 2pjd | 2pjdA01_A |
| 13 | 1g8a | 1g8aA02_A | 53 | 2as0 | 2as0A03_A | 93 | 2pjd | 2pjdA02_A |
| 14 | 1i1n | 1i1nA00_A | 54 | 2avd | 2avdA00_A | 94 | 2plw | 2plwA00_A |
| 15 | 1i4w | 1i4wA01_A | 55 | 2avn | 2avnA00_A | 95 | 2pxx | 2pxxA00_A |
| 16 | 1i9g | 1i9gA02_A | 56 | 2b3t | 2b3tA02_A | 96 | 2py6 | 2py6A03_A |
| 17 | 1inl | 1inlD01_D | 57 | 2b78 | 2b78A03_A | 97 | 2qm3 | 2qm3A02_A |
| 18 | 1jg1 | 1jg1A00_A | 58 | 2b9e | 2b9eA02_A | 98 | 2qrv | 2qrvA01_A |
| 19 | 1jqe | 1jqeA00_A | 59 | 2bm8 | 2bm8B02_B | 99 | 2qy6 | 2qy6A01_A |
| 20 | 1jsx | 1jsxA00_A | 60 | 2cmg | 2cmgA02_A | 100 | 2r3s | 2r3sA03_A |
| 21 | 1l1e | 1l1eB00_B | 61 | 2dpm | 2dpmA01_A | 101 | 2r6z | 2r6zB02_B |
| 22 | 1l3i | 1l3iA00_A | 62 | 2dul | 2dulA01_A | 102 | 2uyo | 2uyoA00_A |
| 23 | 1m6y | 1m6yA01_A | 63 | 2efj | 2efjA02_A | 103 | 2vdw | 2vdwG00_G |
| 24 | 1mjf | 1mjfB02_B | 64 | 2f8l | 2f8lA02_A | 104 | 2vz9 | 2vz9A05_A |
| 25 | 1ne2 | 1ne2B00_B | 65 | 2fhp | 2fhpA00_A | 105 | 2wa2 | 2wa2A00_A |
| 26 | 1nv8 | 1nv8A02_A | 66 | 2fpo | 2fpoC00_C | 106 | 2wa2 | 2wa2A00_B |
| 27 | 1o54 | 1o54A02_A | 67 | 2frx | 2frxB01_B | 107 | 2wk1 | 2wk1A00_A |
| 28 | 1o9g | 1o9gA01_A | 68 | 2g1p | 2g1pB01_B | 108 | 2xvm | 2xvmA00_A |
| 29 | 1p91 | 1p91B00_B | 69 | 2g1p | 2g1pB01_G | 109 | 2xvm | 2xvmA00_B |
| 30 | 1pjz | 1pjzA00_A | 70 | 2gb4 | 2gb4B00_B | 110 | 2xyq | 2xyqA00_A |
| 31 | 1qam | 1qamA01_A | 71 | 2gpy | 2gpyB00_B | 111 | 2yqz | 2yqzA01_A |
| 32 | 1ri5 | 1ri5A00_A | 72 | 2gs9 | 2gs9A01_A | 112 | 2yx1 | 2yx1A03_A |
| 33 | 1u2z | 1u2zA02_A | 73 | 2h00 | 2h00B00_B | 113 | 2yxd | 2yxdA00_A |
| 34 | 1uir | 1uirA02_A | 74 | 2h00 | 2h00B00_C | 114 | 2zfu | 2zfuA02_A |
| 35 | 1uwv | 1uwvA02_A | 75 | 2igt | 2igtA01_A | 115 | 2zig | 2zigA00_A |
| 36 | 1v39 | 1v39A00_A | 76 | 2ih2 | 2ih2A01_A | 116 | 2zwa | 2zwaA01_A |
| 37 | 1vbf | 1vbfA00_A | 77 | 2ih2 | 2ih2A01_B | 117 | 3a27 | 3a27A00_A |
| 38 | 1ve3 | 1ve3A00_A | 78 | 2ih2 | 2ih2A01_C | 118 | 3ajd | 3ajdA02_A |
| 39 | 1vl5 | 1vl5C00_C | 79 | 2ip2 | 2ip2A02_A | 119 | 3b5i | 3b5iB01_B |
| 40 | 1vlm | 1vlmA00_A | 80 | 2jhp | 2jhpA02_A | 120 | 3bkw | 3bkwB00_B |

| No. | PDB | CATH ID | No. | PDB | CATH ID | No. | PDB | CATH ID |
|---|---|---|---|---|---|---|---|---|
| 121 | 3bkx | 3bkxA00_A | 161 | 3g2m | 3g2mA02_A | 201 | 3me5 | 3me5A02_A |
| 122 | 3bt7 | 3bt7A01_A | 162 | 3g5t | 3g5tA00_A | 202 | 3mer | 3merA00_A |
| 123 | 3bt7 | 3bt7A01_C | 163 | 3g7u | 3g7uA01_A | 203 | 3mgg | 3mggB01_B |
| 124 | 3bus | 3busB00_B | 164 | 3g89 | 3g89B00_B | 204 | 3mq2 | 3mq2A00_A |
| 125 | 3bzb | 3bzbB00_B | 165 | 3gdh | 3gdhA00_A | 205 | 3ntv | 3ntvA00_A |
| 126 | 3c3p | 3c3pA00_A | 166 | 3ggd | 3ggdA00_A | 206 | 3o7w | 3o7wA00_A |
| 127 | 3c3y | 3c3yA00_A | 167 | 3giw | 3giwA00_A | 207 | 3ocj | 3ocjA00_A |
| 128 | 3c6k | 3c6kB03_B | 168 | 3gjy | 3gjyA00_A | 208 | 3ofk | 3ofkA00_A |
| 129 | 3cbg | 3cbgA00_A | 169 | 3gnl | 3gnlA01_A | 209 | 3orh | 3orhD00_D |
| 130 | 3cc8 | 3cc8A00_A | 170 | 3gru | 3gruA01_A | 210 | 3ou2 | 3ou2A00_A |
| 131 | 3ccf | 3ccfA00_A | 171 | 3grz | 3grzB00_B | 211 | 3p2e | 3p2eA00_A |
| 132 | 3cgg | 3cggA00_A | 172 | 3gu3 | 3gu3A01_A | 212 | 3p2e | 3p2eA00_B |
| 133 | 3ckk | 3ckkA01_A | 173 | 3gwz | 3gwzA02_A | 213 | 3p9n | 3p9nA00_A |
| 134 | 3cvo | 3cvoA00_A | 174 | 3h2b | 3h2bB00_B | 214 | 3pfg | 3pfgA01_A |
| 135 | 3d2l | 3d2lC01_C | 175 | 3hm2 | 3hm2A00_A | 215 | 3pt9 | 3pt9A02_A |
| 136 | 3dh0 | 3dh0B00_B | 176 | 3hnr | 3hnrA00_A | 216 | 3pta | 3ptaA05_A |
| 137 | 3dlc | 3dlcA00_A | 177 | 3hp7 | 3hp7A02_A | 217 | 3pta | 3ptaA05_B |
| 138 | 3dli | 3dliA00_A | 178 | 3htx | 3htxD03_D | 218 | 3q87 | 3q87B00_B |
| 139 | 3dmg | 3dmgA01_A | 179 | 3htx | 3htxD03_E | 219 | 3qv2 | 3qv2A01_A |
| 140 | 3dmg | 3dmgA02_A | 180 | 3htx | 3htxD03_F | 220 | 3sm3 | 3sm3A00_A |
| 141 | 3dou | 3douA00_A | 181 | 3hvi | 3hviA00_A | 221 | 3ssm | 3ssmC02_C |
| 142 | 3dp7 | 3dp7A03_A | 182 | 3i9f | 3i9fB00_B | 222 | 3thr | 3thrA02_A |
| 143 | 3dr5 | 3dr5A00_A | 183 | 3iht | 3ihtA00_A | 223 | 3thr | 3thrA02_D |
| 144 | 3dtn | 3dtnA01_A | 184 | 3iv6 | 3iv6A01_A | 224 | 3tm4 | 3tm4A02_A |
| 145 | 3duw | 3duwA00_A | 185 | 3iyl | 3iylW04_W | 225 | 3tma | 3tmaA02_A |
| 146 | 3dxy | 3dxyA00_A | 186 | 3jwg | 3jwgA00_A | 226 | 3ua3 | 3ua3A02_A |
| 147 | 3e05 | 3e05B00_B | 187 | 3k0b | 3k0bA02_A | 227 | 3ubt | 3ubtA01_A |
| 148 | 3e23 | 3e23A00_A | 188 | 3k6r | 3k6rA02_A | 228 | 3ubt | 3ubtA01_E |
| 149 | 3e8s | 3e8sA00_A | 189 | 3khk | 3khkA02_A | 229 | 3ubt | 3ubtA01_F |
| 150 | 3eey | 3eeyA00_A | 190 | 3kkz | 3kkzB00_B | 230 | 3ufb | 3ufbA02_A |
| 151 | 3eey | 3eeyA00_D | 191 | 3l8d | 3l8dA00_A | 231 | 3ujc | 3ujcA00_A |
| 152 | 3ege | 3egeA00_A | 192 | 3lcc | 3lccA00_A | 232 | 3uwp | 3uwpA02_A |
| 153 | 3egv | 3egvA02_A | 193 | 3lkd | 3lkdA02_A | 233 | 3uzu | 3uzuA01_A |
| 154 | 3evz | 3evzA01_A | 194 | 3ll7 | 3ll7A02_A | 234 | 3v97 | 3v97A02_A |
| 155 | 3fpf | 3fpfA00_A | 195 | 3lpm | 3lpmA01_A | 235 | 3v97 | 3v97B04_A |
| 156 | 3frh | 3frhA02_A | 196 | 3lst | 3lstA02_A | 236 | 3v97 | 3v97B04_B |
| 157 | 3ftd | 3ftdA01_A | 197 | 3m33 | 3m33A00_A | 237 | 3vyw | 3vywA02_A |
| 158 | 3fut | 3futA01_A | 198 | 3m6u | 3m6uB02_B | 238 | 3x0d | 3x0dA03_A |
| 159 | 3fzg | 3fzgA00_A | 199 | 3mb5 | 3mb5A02_A | 239 | 4a6d | 4a6dA02_A |
| 160 | 3g07 | 3g07A01_A | 200 | 3mcz | 3mczA02_A | 240 | 4auk | 4aukA03_A |

| No. | PDB | CATH ID | No. | PDB | CATH ID |
|---|---|---|---|---|---|
| 241 | 4azs | 4azsA01_A | 281 | 4pne | 4pneA00_A |
| 242 | 4blu | 4bluB00_B | 282 | 4pwy | 4pwyA00_A |
| 243 | 4c08 | 4c08A01_A | 283 | 4qdj | 4qdjA00_A |
| 244 | 4c4a | 4c4aA01_A | 284 | 4qpn | 4qpnA00_A |
| 245 | 4c4a | 4c4aA03_A | 285 | 4qtu | 4qtuB00_B |
| 246 | 4dcm | 4dcmA01_A | 286 | 4qvg | 4qvgA02_A |
| 247 | 4dcm | 4dcmA02_A | 287 | 4rv9 | 4rv9A02_A |
| 248 | 4dkj | 4dkjA01_A | 288 | 4rwz | 4rwzA00_A |
| 249 | 4dkj | 4dkjA01_B | 289 | 4u7t | 4u7tD00_D |
| 250 | 4dkj | 4dkjA01_C | 290 | 4x1o | 4x1oA00_A |
| 251 | 4dmg | 4dmgA03_A | 291 | 4xcx | 4xcxA00_A |
| 252 | 4e2x | 4e2xA02_A | 292 | 4ymh | 4ymhD00_D |
| 253 | 4f85 | 4f85A00_A | 293 | 4yuy | 4yuyB02_B |
| 254 | 4fsd | 4fsdA01_A | 294 | 5c1i | 5c1iA00_A |
| 255 | 4ft4 | 4ft4A02_A | 295 | 5ccb | 5ccbA02_A |
| 256 | 4ft4 | 4ft4A02_B | 296 | 5ciy | 5ciyA01_A |
| 257 | 4fzv | 4fzvA02_A | 297 | 5ciy | 5ciyA01_C |
| 258 | 4gc5 | 4gc5A01_A | 298 | 5ciy | 5ciyA01_D |
| 259 | 4gek | 4gekA00_A | 299 | 5cm2 | 5cm2Z00_Z |
| 260 | 4gqb | 4gqbA02_A | 300 | 5cvd | 5cvdB00_B |
| 261 | 4gua | 4guaA02_A | 301 | 5dm2 | 5dm2A00_A |
| 262 | 4hg2 | 4hg2B01_B | 302 | 5eeh | 5eehC03_C |
| 263 | 4hh4 | 4hh4C01_C | 303 | 5epe | 5epeA00_A |
| 264 | 4htf | 4htfA00_A | 304 | 5ezq | 5ezqA02_A |
| 265 | 4isc | 4iscA00_A | 305 | 5f2k | 5f2kB02_B |
| 266 | 4kdr | 4kdrA00_A | 306 | 5fa8 | 5fa8A00_A |
| 267 | 4kig | 4kigA02_A | 307 | 5fcd | 5fcdA00_A |
| 268 | 4krg | 4krgA01_A | 308 | 5ftw | 5ftwA02_A |
| 269 | 4krg | 4krgA01_B | 309 | 5gm2 | 5gm2K01_K |
| 270 | 4lg1 | 4lg1B00_B | 310 | 5h02 | 5h02A02_A |
| 271 | 4lwo | 4lwoE01_E | 311 | 5hfj | 5hfjC00_C |
| 272 | 4m37 | 4m37A01_A | 312 | 5hoq | 5hoqC00_C |
| 273 | 4m7r | 4m7rA00_A | | | |
| 274 | 4mik | 4mikA00_A | | | |
| 275 | 4mtl | 4mtlA00_A | | | |
| 276 | 4mtl | 4mtlA00_B | | | |
| 277 | 4nec | 4necC01_C | | | |
| 278 | 4obx | 4obxA00_A | | | |
| 279 | 4pca | 4pcaB00_B | | | |
| 280 | 4pio | 4pioB01_B | | | |

**Table S5 List of protein domains used for structural super positions of NAD-binding domains.** 642 structural representatives, corresponding to NAD-binding domains, according to CATH database (Superfamily ID 3.40.50.720).

| No. | PDB | CATH ID | No. | PDB | CATH ID | No. | PDB | CATH ID |
|---|---|---|---|---|---|---|---|---|
| 1 | 1a4iB01_B | a4iB | 41 | 1ks9A01_A | ks9A | 81 | 1t2dA01_A | t2dA |
| 2 | 1bg6A01_A | bg6A | 42 | 1kyqB01_B | kyqB | 82 | 1txgA01_A | txgA |
| 3 | 1bgvA01_A | bgvA | 43 | 1l7dA02_A | l7dA | 83 | 1u8xX01_X | u8xX |
| 4 | 1c0pA01_A | c0pA | 44 | 1lc0A01_A | lc0A | 84 | 1up7A01_A | up7A |
| 5 | 1c1dA02_A | c1dA | 45 | 1lj8A01_A | lj8A | 85 | 1uzmB00_B | uzmB |
| 6 | 1cjcA01_A | cjcA | 46 | 1lqtA01_A | lqtA | 86 | 1vj0A02_A | vj0A |
| 7 | 1cydA00_A | cydA | 47 | 1lqtA01_B | lqtA | 87 | 1vjtA01_A | vjtA |
| 8 | 1dpgA01_A | dpgA | 48 | 1lssA00_A | lssA | 88 | 1vknA01_A | vknA |
| 9 | 1dxyA01_A | dxyA | 49 | 1lu9A02_A | lu9A | 89 | 1vl6B02_B | vl6B |
| 10 | 1dxyA02_A | dxyA | 50 | 1mv8A01_A | mv8A | 90 | 1vpdA01_A | vpdA |
| 11 | 1e6uA01_A | e6uA | 51 | 1mv8A03_A | mv8A | 91 | 1wlyA02_A | wlyA |
| 12 | 1ebfA01_A | ebfA | 52 | 1mx3A01_A | mx3A | 92 | 1wmaA00_A | wmaA |
| 13 | 1edzA01_A | edzA | 53 | 1mx3A02_A | mx3A | 93 | 1wvgA01_A | wvgA |
| 14 | 1eq2A01_A | eq2A | 54 | 1nffA00_A | nffA | 94 | 1x13B01_B | x13B |
| 15 | 1evyA01_A | evyA | 55 | 1nhwA00_A | nhwA | 95 | 1x7dA02_A | x7dA |
| 16 | 1fmcA00_A | fmcA | 56 | 1npyA02_A | npyA | 96 | 1xdwA01_A | xdwA |
| 17 | 1gdhA01_A | gdhA | 57 | 1nytA02_A | nytA | 97 | 1xdwA02_A | xdwA |
| 18 | 1gpjA02_A | gpjA | 58 | 1o0sA04_A | o0sA | 98 | 1xeaA01_A | xeaA |
| 19 | 1gr0A01_A | gr0A | 59 | 1o94A02_A | o94A | 99 | 1xg5B00_B | xg5B |
| 20 | 1gtmA02_A | gtmA | 60 | 1oaaA00_A | oaaA | 100 | 1xgkA01_A | xgkA |
| 21 | 1gu7A02_A | gu7A | 61 | 1oi7A01_A | oi7A | 101 | 1xq1A00_A | xq1A |
| 22 | 1h2bA02_A | h2bA | 62 | 1omoA02_A | omoA | 102 | 1xq6A00_A | xq6A |
| 23 | 1h5qA00_A | h5qA | 63 | 1ooeA00_A | ooeA | 103 | 1xu9C00_C | xu9C |
| 24 | 1hdoA00_A | hdoA | 64 | 1orrC00_C | orrC | 104 | 1y1pA01_A | y1pA |
| 25 | 1hyeA01_A | hyeA | 65 | 1p1hB02_B | p1hB | 105 | 1y81A00_A | y81A |
| 26 | 1hyhA01_A | hyhA | 66 | 1p1jB01_B | p1jB | 106 | 1y8qB01_B | y8qB |
| 27 | 1i24A01_A | i24A | 67 | 1p3dA01_A | p3dA | 107 | 1y8qC00_C | y8qC |
| 28 | 1i36A01_A | i36A | 68 | 1pj3A02_A | pj3A | 108 | 1y8qC00_D | y8qC |
| 29 | 1id1A00_A | id1A | 69 | 1pjqA01_A | pjqA | 109 | 1yb1B01_B | yb1B |
| 30 | 1isiA02_A | isiA | 70 | 1pl8A02_A | pl8A | 110 | 1yb5A02_A | yb5A |
| 31 | 1iukA00_A | iukA | 71 | 1ps9A02_A | ps9A | 111 | 1yo6F00_F | yo6F |
| 32 | 1iz0A02_A | iz0A | 72 | 1q0qA01_A | q0qA | 112 | 1yovC01_C | yovC |
| 33 | 1j4aD01_D | j4aD | 73 | 1qp8A01_A | qp8A | 113 | 1yqgA01_A | yqgA |
| 34 | 1j6uA01_A | j6uA | 74 | 1qp8A02_A | qp8A | 114 | 1yrlB01_B | yrlB |
| 35 | 1jayA00_A | jayA | 75 | 1r12A02_A | r12A | 115 | 1yxmC00_C | yxmC |
| 36 | 1jtvA00_A | jtvA | 76 | 1r6dA01_A | r6dA | 116 | 1z45A01_A | z45A |
| 37 | 1jvbA02_A | jvbA | 77 | 1sbyA00_A | sbyA | 117 | 1z82A01_A | z82A |
| 38 | 1jw9B00_B | jw9B | 78 | 1sc6A01_A | sc6A | 118 | 1zcjA02_A | zcjA |
| 39 | 1kc0A00_A | kc0A | 79 | 1snyA00_A | snyA | 119 | 1zejA01_A | zejA |
| 40 | 1kolA02_A | kolA | 80 | 1spxA00_A | spxA | 120 | 1zemA00_A | zemA |

| No. | PDB | CATH ID | No. | PDB | CATH ID | No. | PDB | CATH ID |
|---|---|---|---|---|---|---|---|---|
| 121 | 1zh8A01_A | zh8A | 161 | 2g0tB01_B | g0tB | 201 | 2qrlA01_A | qrlA |
| 122 | 1zk4A00_A | zk4A | 162 | 2g1uA00_A | g1uA | 202 | 2qytA01_A | qytA |
| 123 | 2a35A00_A | a35A | 163 | 2g5cA01_A | g5cA | 203 | 2r6jA01_A | r6jA |
| 124 | 2a4kB01_B | a4kB | 164 | 2g76A01_A | g76A | 204 | 2rafB01_B | rafB |
| 125 | 2aefA01_A | aefA | 165 | 2g76A02_A | g76A | 205 | 2rcyA01_A | rcyA |
| 126 | 2amfA01_A | amfA | 166 | 2g82A01_A | g82A | 206 | 2rhcA00_A | rhcA |
| 127 | 2amfA01_D | amfA | 167 | 2gcgA02_A | gcgA | 207 | 2uv8A04_A | uv8A |
| 128 | 2axqA01_A | axqA | 168 | 2gdzA00_A | gdzA | 208 | 2vn8A02_A | vn8A |
| 129 | 2b0jA01_A | b0jA | 169 | 2ggsA01_A | ggsA | 209 | 2vq3A00_A | vq3A |
| 130 | 2b4qB00_B | b4qB | 170 | 2h6eA02_A | h6eA | 210 | 2w2kA01_A | w2kA |
| 131 | 2b5wA02_A | b5wA | 171 | 2hjsA01_A | hjsA | 211 | 2w2kA02_A | w2kA |
| 132 | 2b69A01_A | b69A | 172 | 2hmtA00_A | hmtA | 212 | 2we8A02_A | we8A |
| 133 | 2bd0A01_A | bd0A | 173 | 2hrzA01_A | hrzA | 213 | 2wm3A01_A | wm3A |
| 134 | 2bgkA00_A | bgkA | 174 | 2i76A01_A | i76A | 214 | 2x4gA00_A | x4gA |
| 135 | 2bh9A01_A | bh9A | 175 | 2ixaA01_A | ixaA | 215 | 2x5oA01_A | x5oA |
| 136 | 2bi7A01_A | bi7A | 176 | 2izzB01_B | izzB | 216 | 2x9gD00_D | x9gD |
| 137 | 2bkaA00_A | bkaA | 177 | 2j8zA02_A | j8zA | 217 | 2y1eA01_A | y1eA |
| 138 | 2bmaA02_A | bmaA | 178 | 2jahA00_A | jahA | 218 | 2ydyA00_A | ydyA |
| 139 | 2c0cA02_A | c0cA | 179 | 2jhfA02_A | jhfA | 219 | 2yutA00_A | yutA |
| 140 | 2c2xA01_A | c2xA | 180 | 2jl1A01_A | jl1A | 220 | 2yyyA01_A | yyyA |
| 141 | 2c5aA01_A | c5aA | 181 | 2nqtA01_A | nqtA | 221 | 2z1nA00_A | z1nA |
| 142 | 2cdcA02_A | cdcA | 182 | 2nvwB01_B | nvwB | 222 | 2zatA00_A | zatA |
| 143 | 2cukA01_A | cukA | 183 | 2o23A00_A | o23A | 223 | 2zb4A02_A | zb4A |
| 144 | 2d0iA01_A | d0iA | 184 | 2o3jA03_A | o3jA | 224 | 2ztuB00_B | ztuB |
| 145 | 2d1yA00_A | d1yA | 185 | 2o3jB01_B | o3jB | 225 | 3a06B01_B | a06B |
| 146 | 2d4aD01_D | d4aD | 186 | 2o3sA02_A | o3sA | 226 | 3abiA01_A | abiA |
| 147 | 2d59A00_A | d59A | 187 | 2o4cA01_A | o4cA | 227 | 3adoA01_A | adoA |
| 148 | 2d8aA02_A | d8aA | 188 | 2o4cA02_A | o4cA | 228 | 3afmB00_B | afmB |
| 149 | 2dc1A01_A | dc1A | 189 | 2o7sA03_A | o7sA | 229 | 3aoeC03_C | aoeC |
| 150 | 2dknB00_B | dknB | 190 | 2obnD01_D | obnD | 230 | 3asuB01_B | asuB |
| 151 | 2dtxA00_A | dtxA | 191 | 2p2sA01_A | p2sA | 231 | 3b1fA01_A | b1fA |
| 152 | 2dvmB02_B | dvmB | 192 | 2p4hX00_X | p4hX | 232 | 3ba1A01_A | ba1A |
| 153 | 2eggA02_A | eggA | 193 | 2ph5A01_A | ph5A | 233 | 3ba1A02_A | ba1A |
| 154 | 2ejwA01_A | ejwA | 194 | 2pk3A01_A | pk3A | 234 | 3c1aA01_A | c1aA |
| 155 | 2eklA01_A | eklA | 195 | 2pv7A01_A | pv7A | 235 | 3c7aA01_A | c7aA |
| 156 | 2eklA02_A | eklA | 196 | 2py6A02_A | py6A | 236 | 3c85A00_A | c85A |
| 157 | 2et6A02_A | et6A | 197 | 2q1sA01_A | q1sA | 237 | 3c8mA01_A | c8mA |
| 158 | 2ew8B00_B | ew8B | 198 | 2q1wC00_C | q1wC | 238 | 3cinA01_A | cinA |
| 159 | 2f1kA01_A | f1kA | 199 | 2qq5A00_A | qq5A | 239 | 3d1lB01_B | d1lB |
| 160 | 2fwmX00_X | fwmX | 200 | 2qrjA02_A | qrjA | 240 | 3d4oA02_A | d4oA |

| No. | PDB | CATH ID | No. | PDB | CATH ID | No. | PDB | CATH ID |
|---|---|---|---|---|---|---|---|---|
| 241 | 3d4oB01_B | d4oB | 281 | 3gg2C03_C | gg2C | 321 | 3k5pA02_A | k5pA |
| 242 | 3d7lA00_A | d7lA | 282 | 3gg2C03_D | gg2C | 322 | 3k6jA01_A | k6jA |
| 243 | 3db2A01_A | db2A | 283 | 3gg9A01_A | gg9A | 323 | 3k96A01_A | k96A |
| 244 | 3dfuA01_A | dfuA | 284 | 3gg9A02_A | gg9A | 324 | 3kb6A01_A | kb6A |
| 245 | 3dfzB01_B | dfzB | 285 | 3ghyA01_A | ghyA | 325 | 3keoB02_B | keoB |
| 246 | 3dl2A01_A | dl2A | 286 | 3gmsA02_A | gmsA | 326 | 3ko8A01_A | ko8A |
| 247 | 3dmyA01_A | dmyA | 287 | 3gohA02_A | gohA | 327 | 3ktdC01_C | ktdC |
| 248 | 3do5A01_A | do5A | 288 | 3gpiA00_A | gpiA | 328 | 3kzvA00_A | kzvA |
| 249 | 3dqpA00_A | dqpA | 289 | 3gqvA02_A | gqvA | 329 | 3l4bC01_A | l4bC |
| 250 | 3dr3A01_A | dr3A | 290 | 3gt0A01_A | gt0A | 330 | 3l4bC01_C | l4bC |
| 251 | 3dttA00_A | dttA | 291 | 3guyA00_A | guyA | 331 | 3l6dA01_A | l6dA |
| 252 | 3dtyA01_A | dtyA | 292 | 3gvxA02_A | gvxA | 332 | 3l77A00_A | l77A |
| 253 | 3e03A00_A | e03A | 293 | 3gvxB01_B | gvxB | 333 | 3l9wA01_A | l9wA |
| 254 | 3e18A01_A | e18A | 294 | 3h2sA00_A | h2sA | 334 | 3l9wA01_B | l9wA |
| 255 | 3e48A01_A | e48A | 295 | 3h2zA01_A | h2zA | 335 | 3lf2A01_A | lf2A |
| 256 | 3e82B01_B | e82B | 296 | 3h5nD02_C | h5nD | 336 | 3lk7A01_A | lk7A |
| 257 | 3e8xA00_A | e8xA | 297 | 3h5nD02_D | h5nD | 337 | 3llvA00_A | llvA |
| 258 | 3e9nA00_A | e9nA | 298 | 3h7aA00_A | h7aA | 338 | 3lvfP01_P | lvfP |
| 259 | 3egoA01_A | egoA | 299 | 3h8vB00_B | h8vB | 339 | 3m1aJ00_J | m1aJ |
| 260 | 3evtA01_A | evtA | 300 | 3hdjA02_A | hdjA | 340 | 3m2pB00_B | m2pB |
| 261 | 3ezyA01_A | ezyA | 301 | 3hg7A01_A | hg7A | 341 | 3m2tB01_B | m2tB |
| 262 | 3f1lB00_B | f1lB | 302 | 3hg7A02_A | hg7A | 342 | 3m6iA02_A | m6iA |
| 263 | 3f4lA01_A | f4lA | 303 | 3hhpA01_A | hhpA | 343 | 3moiA01_A | moiA |
| 264 | 3fbgA02_A | fbgA | 304 | 3hn2B01_B | hn2B | 344 | 3mtjA01_A | mtjA |
| 265 | 3ff4A00_A | ff4A | 305 | 3hn7A01_A | hn7A | 345 | 3mw9A03_A | mw9A |
| 266 | 3fhlA01_A | fhlA | 306 | 3hwrA01_A | hwrA | 346 | 3mweB01_B | mweB |
| 267 | 3fi9A01_A | fi9A | 307 | 3i4fC00_C | i4fC | 347 | 3n74A00_A | n74A |
| 268 | 3fpcA02_A | fpcA | 308 | 3i83A01_A | i83A | 348 | 3nklB00_B | nklB |
| 269 | 3fr7A01_A | fr7A | 309 | 3ic5A00_A | ic5A | 349 | 3nv9A02_A | nv9A |
| 270 | 3fwzA00_A | fwzA | 310 | 3idsA01_A | idsA | 350 | 3nx4A02_A | nx4A |
| 271 | 3fwzA00_B | fwzA | 311 | 3imfC00_C | imfC | 351 | 3nywD00_D | nywD |
| 272 | 3g0oA01_A | g0oA | 312 | 3ip1A02_A | ip1A | 352 | 3o38B01_B | o38B |
| 273 | 3g79A01_A | g79A | 313 | 3ip3A01_A | ip3A | 353 | 3o8qA02_A | o8qA |
| 274 | 3g79A02_A | g79A | 314 | 3is3A00_A | is3A | 354 | 3o9zA01_A | o9zA |
| 275 | 3gazA02_A | gazA | 315 | 3iupA02_A | iupA | 355 | 3oh8A02_A | oh8A |
| 276 | 3gedA00_A | gedA | 316 | 3iusB00_B | iusB | 356 | 3ohsX01_X | ohsX |
| 277 | 3gemD00_D | gemD | 317 | 3jtmA01_A | jtmA | 357 | 3oidC00_C | oidC |
| 278 | 3gg2A01_A | gg2A | 318 | 3jtmA02_A | jtmA | 358 | 3oigA00_A | oigA |
| 279 | 3gg2A01_D | gg2A | 319 | 3jv7A02_A | jv7A | 359 | 3oj0A00_A | oj0A |
| 280 | 3gg2C03_A | gg2C | 320 | 3jyoA02_A | jyoA | 360 | 3omlA01_A | omlA |

| No. | PDB | CATH ID | No. | PDB | CATH ID | No. | PDB | CATH ID |
|---|---|---|---|---|---|---|---|---|
| 361 | 3on5B02_B | on5B | 401 | 3tnlA02_A | tnlA | 441 | 4c4oA02_A | c4oA |
| 362 | 3ondA02_A | ondA | 402 | 3triA01_A | triA | 442 | 4cpdA02_A | cpdA |
| 363 | 3op4B00_B | op4B | 403 | 3u0bA01_A | u0bA | 443 | 4cr6A00_A | cr6A |
| 364 | 3oqbA01_A | oqbA | 404 | 3u49D00_D | u49D | 444 | 4cujA01_A | cujA |
| 365 | 3pduA01_A | pduA | 405 | 3u62A02_A | u62A | 445 | 4cujA02_A | cujA |
| 366 | 3phhA02_A | phhA | 406 | 3u9lA00_A | u9lA | 446 | 4d79A00_A | d79A |
| 367 | 3pi7A02_A | pi7A | 407 | 3uceA00_A | uceA | 447 | 4dllA01_A | dllA |
| 368 | 3pidA01_A | pidA | 408 | 3ucxA00_A | ucxA | 448 | 4dplA01_A | dplA |
| 369 | 3pidA03_A | pidA | 409 | 3ulkA01_A | ulkA | 449 | 4dyvA00_A | dyvA |
| 370 | 3pk0D00_D | pk0D | 410 | 3un1C00_C | un1C | 450 | 4e12A01_A | e12A |
| 371 | 3plnA03_A | plnA | 411 | 3uogA02_A | uogA | 451 | 4e21A01_A | e21A |
| 372 | 3pvzD01_D | pvzD | 412 | 3uuwA01_A | uuwA | 452 | 4e2xA04_A | e2xA |
| 373 | 3pwzA02_A | pwzA | 413 | 3uw3B01_B | uw3B | 453 | 4e3zB00_B | e3zB |
| 374 | 3q2iA01_A | q2iA | 414 | 3v2uA01_A | v2uA | 454 | 4e4yA00_A | e4yA |
| 375 | 3qhaA01_A | qhaA | 415 | 3v8bC00_C | v8bC | 455 | 4e5nA02_A | e5nA |
| 376 | 3qivA00_A | qivA | 416 | 3vpsB01_B | vpsB | 456 | 4e5nC01_C | e5nC |
| 377 | 3qsgA01_A | qsgA | 417 | 3vtfA02_A | vtfA | 457 | 4e5yC01_C | e5yC |
| 378 | 3qvoA00_A | qvoA | 418 | 3vvcA01_A | vvcA | 458 | 4ej6A02_A | ej6A |
| 379 | 3qvsA01_A | qvsA | 419 | 3wb9A01_A | wb9A | 459 | 4ew6A01_A | ew6A |
| 380 | 3qy9B01_B | qy9B | 420 | 3wfiA01_A | wfiA | 460 | 4eyeA02_A | eyeA |
| 381 | 3r1iB00_B | r1iB | 421 | 3wfjH01_H | wfjH | 461 | 4f6cB00_B | f6cB |
| 382 | 3r3sA00_A | r3sA | 422 | 3wg9A02_A | wg9A | 462 | 4fc7D00_D | fc7D |
| 383 | 3r6dA00_A | r6dA | 423 | 3wgtA01_A | wgtA | 463 | 4fdaA00_A | fdaA |
| 384 | 3rc1A01_A | rc1A | 424 | 3widA02_A | widA | 464 | 4fflA01_A | fflA |
| 385 | 3rd5A00_A | rd5A | 425 | 3wj7A00_A | wj7A | 465 | 4fgwA01_A | fgwA |
| 386 | 3rfxA01_A | rfxA | 426 | 3wnvA01_A | wnvA | 466 | 4g2nA01_A | g2nA |
| 387 | 3rkrA00_A | rkrA | 427 | 3ws7A01_A | ws7A | 467 | 4g2nA02_A | g2nA |
| 388 | 3rufA01_A | rufA | 428 | 3wyeB00_B | wyeB | 468 | 4g65A01_A | g65A |
| 389 | 3ruiA00_A | ruiA | 429 | 3zv4A00_A | zv4A | 469 | 4g65A03_A | g65A |
| 390 | 3rwbA00_A | rwbA | 430 | 4a0sA02_A | a0sA | 470 | 4gbjC01_C | gbjC |
| 391 | 3s2eA02_A | s2eA | 431 | 4a0sA02_D | a0sA | 471 | 4gh5B00_B | gh5B |
| 392 | 3s8mA00_A | s8mA | 432 | 4a27A02_A | a27A | 472 | 4gkbA00_A | gkbA |
| 393 | 3slgF01_F | slgF | 433 | 4b3hA02_A | b3hA | 473 | 4gwgA01_A | gwgA |
| 394 | 3svtA00_A | svtA | 434 | 4b4oG00_G | b4oG | 474 | 4gx0A02_A | gx0A |
| 395 | 3sx2H00_H | sx2H | 435 | 4bguA01_A | bguA | 475 | 4gx0B04_B | gx0B |
| 396 | 3sxpA01_A | sxpA | 436 | 4bmnD00_D | bmnD | 476 | 4h15A00_A | h15A |
| 397 | 3t4xA00_A | t4xA | 437 | 4bmvI00_I | bmvI | 477 | 4h3vA01_A | h3vA |
| 398 | 3tfoB00_B | tfoB | 438 | 4bs9A02_A | bs9A | 478 | 4h7pA01_A | h7pA |
| 399 | 3tjrA00_A | tjrA | 439 | 4bucA01_A | bucA | 479 | 4hadA01_A | hadA |
| 400 | 3tl2A01_A | tl2A | 440 | 4bvaA02_A | bvaA | 480 | 4hfmA02_A | hfmA |

| No. | PDB | CATH ID | No. | PDB | CATH ID | No. | PDB | CATH ID |
|---|---|---|---|---|---|---|---|---|
| 481 | 4hktA01_A | hktA | 521 | 4n18A01_A | n18A | 561 | 4x54A00_A | x54A |
| 482 | 4hp8A00_A | hp8A | 522 | 4n5mA00_A | n5mA | 562 | 4xa8A01_A | xa8A |
| 483 | 4hujA00_A | hujA | 523 | 4n7rA02_A | n7rA | 563 | 4xb1A01_A | xb1A |
| 484 | 4hxyA00_A | hxyA | 524 | 4nbuA00_A | nbuA | 564 | 4xcvA01_A | xcvA |
| 485 | 4hy3A02_A | hy3A | 525 | 4nheB01_B | nheB | 565 | 4xcvA02_A | xcvA |
| 486 | 4hy3D01_D | hy3D | 526 | 4nimA00_A | nimA | 566 | 4xgiA03_A | xgiA |
| 487 | 4id9A01_A | id9A | 527 | 4njmA01_A | njmA | 567 | 4xijA02_A | xijA |
| 488 | 4idcA02_A | idcA | 528 | 4njmA02_A | njmA | 568 | 4xqcA01_A | xqcA |
| 489 | 4ii2A04_A | ii2A | 529 | 4o6vA00_A | o6vA | 569 | 4xr9B01_B | xr9B |
| 490 | 4ilkA02_A | ilkA | 530 | 4oaqA02_A | oaqA | 570 | 4xr9B02_A | xr9B |
| 491 | 4impA03_A | impA | 531 | 4oh1A02_A | oh1A | 571 | 4xr9B02_B | xr9B |
| 492 | 4imrB00_B | imrB | 532 | 4okiA00_A | okiA | 572 | 4xybA01_A | xybA |
| 493 | 4inaA01_A | inaA | 533 | 4ol9A01_A | ol9A | 573 | 4xymA01_A | xymA |
| 494 | 4is2A00_A | is2A | 534 | 4om8A01_A | om8A | 574 | 4yaeA00_A | yaeA |
| 495 | 4iuyA00_A | iuyA | 535 | 4oneC00_C | oneC | 575 | 4ycaB01_B | ycaB |
| 496 | 4izhA02_A | izhA | 536 | 4oo3A01_A | oo3A | 576 | 4ydrA01_A | ydrA |
| 497 | 4j0eA01_A | j0eA | 537 | 4p22A01_A | p22A | 577 | 4ypoA01_A | ypoA |
| 498 | 4j1qA00_A | j1qA | 538 | 4pg7A01_A | pg7A | 578 | 4yqyB00_B | yqyB |
| 499 | 4j4hA02_A | j4hA | 539 | 4prkA01_A | prkA | 579 | 4yqzB00_B | yqzB |
| 500 | 4j6fA02_A | j6fA | 540 | 4pvdD00_D | pvdD | 580 | 4yrbA00_A | yrbA |
| 501 | 4jbgA02_A | jbgA | 541 | 4q9nA00_A | q9nA | 581 | 4yrbF00_F | yrbF |
| 502 | 4jgbB00_B | jgbB | 542 | 4qedA00_A | qedA | 582 | 4yrdA01_A | yrdA |
| 503 | 4jp2A00_A | jp2A | 543 | 4qqrB00_B | qqrB | 583 | 4yt2A01_A | yt2A |
| 504 | 4k28A02_A | k28A | 544 | 4r01A00_A | r01A | 584 | 4ywjA01_A | ywjA |
| 505 | 4koaA01_A | koaA | 545 | 4r16A01_A | r16A | 585 | 4z0tA00_A | z0tA |
| 506 | 4kzpB00_B | kzpB | 546 | 4r16A02_A | r16A | 586 | 4z9fA00_A | z9fA |
| 507 | 4lgvA01_A | lgvA | 547 | 4r16A02_B | r16A | 587 | 4zd6C00_C | zd6C |
| 508 | 4lmpA01_A | lmpA | 548 | 4r1sB00_B | r1sB | 588 | 4zgwB00_B | zgwB |
| 509 | 4lmpA02_A | lmpA | 549 | 4r3nA01_A | r3nA | 589 | 4zjuA00_A | zjuA |
| 510 | 4lrtB01_B | lrtB | 550 | 4s1vB01_B | s1vB | 590 | 4zqbB01_B | zqbB |
| 511 | 4lswA01_A | lswA | 551 | 4tqgA00_A | tqgA | 591 | 4zrmA01_A | zrmA |
| 512 | 4lvuA00_A | lvuA | 552 | 4u5qB00_B | u5qB | 592 | 5a04A01_A | a04A |
| 513 | 4lw8A01_A | lw8A | 553 | 4uejA02_A | uejA | 593 | 5a3vA02_A | a3vA |
| 514 | 4m1qA01_A | m1qA | 554 | 4urfA00_A | urfA | 594 | 5a9tA01_A | a9tA |
| 515 | 4m55E00_E | m55E | 555 | 4w4tB00_B | w4tB | 595 | 5aovA01_A | aovA |
| 516 | 4m55F00_F | m55F | 556 | 4w6zA02_A | w6zA | 596 | 5aovA02_A | aovA |
| 517 | 4miyA01_A | miyA | 557 | 4wecC00_C | wecC | 597 | 5ayvA01_A | ayvA |
| 518 | 4mkxA01_A | mkxA | 558 | 4wkgA02_A | wkgA | 598 | 5bseA01_A | bseA |
| 519 | 4mowC00_C | mowC | 559 | 4wpgA01_A | wpgA | 599 | 5bt9D00_D | bt9D |
| 520 | 4mp8A02_A | mp8A | 560 | 4wuvA00_A | wuvA | 600 | 5c37C02_C | c37C |

| No. | PDB | CATH ID | No. | PDB | CATH ID | No. | PDB | CATH ID |
|---|---|---|---|---|---|---|---|---|
| 601 | 5dcyB00_B | dcyB | 641 | 5u4qB00_B | u4qB | | | |
| 602 | 5dp2A02_A | dp2A | 642 | 5u9cA01_A | u9cA | | | |
| 603 | 5dzsB02_B | dzsB | | | | | | |
| 604 | 5eesA01_A | eesA | | | | | | |
| 605 | 5epoA00_A | epoA | | | | | | |
| 606 | 5er6C00_C | er6C | | | | | | |
| 607 | 5er9B01_B | er9B | | | | | | |
| 608 | 5f5nA00_A | f5nA | | | | | | |
| 609 | 5ff5B02_B | ff5B | | | | | | |
| 610 | 5fi3A02_A | fi3A | | | | | | |
| 611 | 5fydB00_B | fydB | | | | | | |
| 612 | 5g0tC00_C | g0tC | | | | | | |
| 613 | 5g4kA00_A | g4kA | | | | | | |
| 614 | 5gy7A01_A | gy7A | | | | | | |
| 615 | 5gz3A01_A | gz3A | | | | | | |
| 616 | 5idqB00_B | idqB | | | | | | |
| 617 | 5if3B00_B | if3B | | | | | | |
| 618 | 5ilgB00_B | ilgB | | | | | | |
| 619 | 5in4B01_B | in4B | | | | | | |
| 620 | 5itwA00_A | itwA | | | | | | |
| 621 | 5iz4A00_A | iz4A | | | | | | |
| 622 | 5jazA01_A | jazA | | | | | | |
| 623 | 5je8B01_B | je8B | | | | | | |
| 624 | 5jlaA00_A | jlaA | | | | | | |
| 625 | 5jo9A00_A | jo9A | | | | | | |
| 626 | 5jy1A00_A | jy1A | | | | | | |
| 627 | 5kkcD01_D | kkcD | | | | | | |
| 628 | 5ktkA00_A | ktkA | | | | | | |
| 629 | 5kvcA00_A | kvcA | | | | | | |
| 630 | 5kvsB01_B | kvsB | | | | | | |
| 631 | 5l4lA00_A | l4lA | | | | | | |
| 632 | 5l9aB00_B | l9aB | | | | | | |
| 633 | 5ldgA00_A | ldgA | | | | | | |
| 634 | 5t57A01_A | t57A | | | | | | |
| 635 | 5t5qB00_B | t5qB | | | | | | |
| 636 | 5thkA00_A | thkA | | | | | | |
| 637 | 5thqA00_A | thqA | | | | | | |
| 638 | 5tnxA02_A | tnxA | | | | | | |
| 639 | 5tt0B00_B | tt0B | | | | | | |
| 640 | 5tx7A01_A | tx7A | | | | | | |
